# When mitochondria fall apart: Unbalanced mitochondrial segregation triggers loss of mtDNA in the absence of mitochondrial fusion

**DOI:** 10.1101/2025.05.13.653688

**Authors:** Lisa Dengler, Francesco Padovani, Bianca Lemke, Rebecca Brugger, Benedikt Westermann, Boris Maček, Kurt M. Schmoller, Jennifer C. Ewald

## Abstract

Mitochondrial biogenesis and inheritance must be carefully regulated alongside cell division to ensure proper mitochondrial function and cell survival. The dynamics of the mitochondrial network, including fusion and fission, play a crucial role in mitochondrial inheritance by facilitating the distribution and quality control of mitochondria. In budding yeast, simultaneous inhibition of both fusion and fission leads to loss of mitochondrial DNA (mtDNA) integrity, resulting in an increased frequency of petite cells. Loss of mitochondrial fusion alone results in the complete loss of mtDNA. While the loss of mtDNA in the absence of mitochondrial fusion has been known for almost 30 years, the reason remained unclear. Here, we investigate the consequences of impaired mitochondrial fusion through depletion of the mitofusin Fzo1. We follow the emerging phenotype by live-cell imaging and the analysis of more than thirty thousand single cells across their cell cycle. Fzo1 depletion causes rapid mitochondrial fragmentation and a reduction in mitochondrial membrane potential, followed by a progressive decline of mtDNA content and cellular growth rate over several cell divisions. During division, Fzo1-depleted daughters obtain an unusually large amount of mitochondria, leaving the mother with too little. This results in a strong disbalance of mitochondrial mass in the population. Additionally, Fzo1-depleted cells lose the ability to adjust mtDNA synthesis to compensate for a low mitochondrial content. The combined effects of unequal distribution and reduced synthesis drive rapid mtDNA loss. These results show how fusion defects lead to mtDNA loss and mitochondrial dysfunction, contributing to understanding diseases linked to fusion defects.

## Introduction

Mitochondrial biogenesis needs to be tightly coordinated with cell proliferation. Since mitochondria cannot be formed de novo, their proper inheritance during the cell division cycle is essential for cell survival. The accurate distribution and quality control during cell division is enabled by the dynamics of the mitochondrial network, including fusion and fission. Mitochondrial fusion was suggested to increase content mixing within the mitochondrial network, while fission helps to remove non-functional mitochondria through mitophagy (Abeliovich, 2023; Friedman & Nunnari, 2014; Khan et al., 2024; Quintana-Cabrera & Scorrano, 2023; Westermann, 2010). In humans, impaired mitochondrial fusion leads to a strong reduction of mitochondrial DNA (mtDNA). Accordingly, fusion defects are associated with numerous diseases, such as peripheral neuropathy and optic atrophy, which are frequently connected to the depletion of mtDNA (Alexander et al., 2000; Delettre et al., 2000; Züchner et al., 2004). However, how and why the loss of fusion leads to a depletion of mtDNA is still not understood.

Unlike human cells, the budding yeast *S. cerevisiae* can tolerate the complete loss of mtDNA, making it an ideal system to unravel the connection between fusion, fission and mtDNA maintenance. The budding yeast mtDNA encodes four subunits of the respiratory chain, three subunits of the ATP synthase (complex V) and one subunit of the mitochondrial ribosome. Loss of mtDNA not only leads to the loss of mtDNA encoded subunits of the respiratory chain but also a strong reduction of the nuclear encoded respiratory chain subunits (Dagsgaard et al., 2001; Vowinckel et al., 2021). Respiration is involved in maintaining the proton gradient across the inner membrane, which is a major part of the mitochondrial membrane potential (MMP). Maintaining a high MMP is required for mitochondrial import of nuclear encoded mitochondrial proteins, which are crucial for mitochondrial biogenesis and cellular health. Thus, while loss of mtDNA is not lethal to yeast, its loss results in the inability to respire and reduced cellular growth, resulting in the so-called “petite phenotype”.

An increased frequency of petite cells is observed when fusion and fission of mitochondria are simultaneously inhibited, which likely leads to defects in mitochondrial DNA (mtDNA) integrity (Osman et al., 2015; Wisniewski et al., 2024). Loss of mitochondrial fission alone only mildly increases petite frequency, whereas the absence of fusion has severe effects: Cells lacking the mitofusin Fzo1, a GTPase required for fusion of the mitochondrial outer membrane, show complete loss of mtDNA in the entire population. Even though this effect has been known for almost 30 years (Hermann et al., 1998; Rapaport et al., 1998), it is still unclear why the fragmentation of mitochondria lacking Fzo1 causes the loss of their genome. Exploring the cause has been hampered by the many defects of the deletion mutant, including fragmented mitochondria, a reduced MMP and slow growth. To disentangle primary causes and secondary effects, it is necessary to monitor cells immediately as the phenotype emerges.

Here, we revealed the dynamics of mtDNA loss in fusion-deficient cells by controlled depletion of Fzo1. We analyzed mitochondrial morphology, distribution and function in ten thousands of dividing cells using live-cell imaging. While the primary phenotype, the fragmentation of mitochondria, is established within less than an hour, the loss of mtDNA and reduction of growth manifested over 21 hours (10 generations). We found that despite an initial delay of mitochondrial transport to the bud, at the end of the cell cycle, an unusually large amount of mitochondria is transported into the bud, resulting in mother cells with low mitochondrial concentration and mtDNA content. Cells with low mitochondrial content have a reduced ability to synthesize mtDNA encoded proteins, resulting in the rapid development of the Δ*fzo1* phenotype.

## Results

### Loss of Fzo1 leads to rapid changes in mitochondrial morphology followed by a gradual decline in growth rate

Complex phenotypes of mutants are difficult to study, if they are characterized by multiple interdependent defects as in the Δ*fzo1* mutant. The cause-and-effect relationships between the individual defects are often not clearly understood and therefore the trigger of the phenotype is often unclear. To unravel the primary causes of such complex phenotypes, a system is required that removes the protein of interest specifically and rapidly. Here, we use the AID (auxin-induced degron) system, which enables induced protein degradation within a short time frame without additional perturbations to the cell (Yesbolatova et al., 2020). In the presence of auxin, the TIR protein, an adapter for E3 ubiquitin ligases, induces the degradation of AID-tagged proteins.

Since we observed background degradation of Fzo1 using constitutively expressed TIR protein, we expressed TIR under an inducible promoter (using either anhydrotetracycline (aTC) or β- estradiol, see Table S1 for strains) shortly before depletion. Using this system, depletion of FLAG-AID-Fzo1 is achieved within 5-15 min and no background degradation is observed before depletion (Fig 1A).

**Figure 1.**
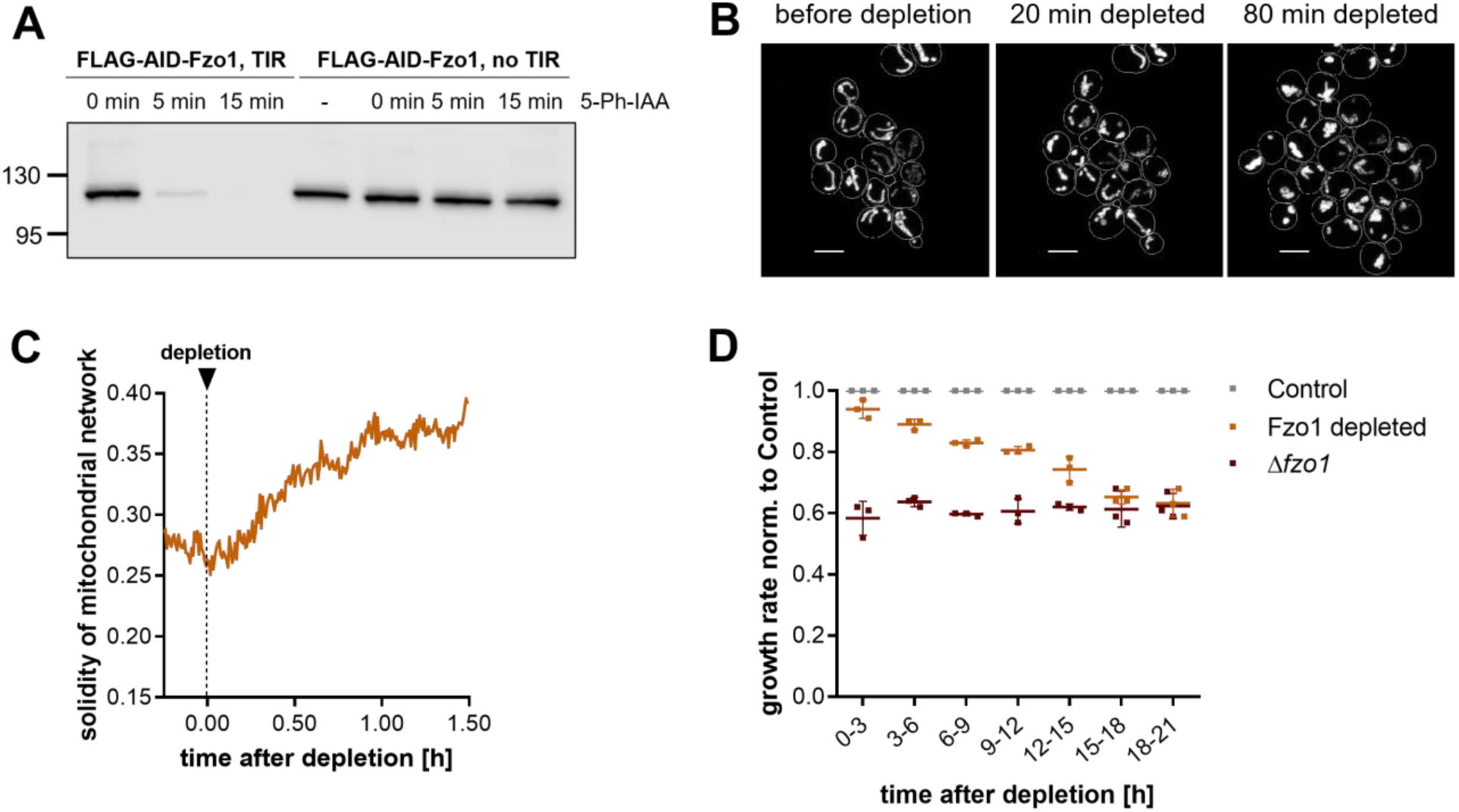
Loss of Fzo1 leads to rapid changes in mitochondrial morphology. Cells were grown in synthetic complete (SC) medium containing 1 % glucose and Fzo1 depletion was induced at the indicated times. A) Western blot analysis of FLAG-AID-Fzo1. TIR expression was induced by addition of anhydrotetracycline (aTC) 2 h before depletion. Then, Fzo1 depletion was induced by addition of 5-Ph-IAA at t = 0 min. A representative blot from two biological replicates is shown. B) Example pictures of confocal microscopy of mitochondria. Maximum z-projections of mitochondria visualized by preSu9-mCardinal before and after depletion of Fzo1 are shown. Scale bar = 5 µm. C) Solidity of the mitochondrial network after Fzo1 depletion. Mean of two biological replicates with a total of 200 cells. D) Growth rates determined in three-hour intervals, normalized to the Control at each time interval. Dots indicate data from three biological replicates, bars indicate mean and standard deviation.

Since depletion is achieved rapidly and completely, we can combine this system with microfluidics-based live-cell imaging to precisely determine how rapidly loss of Fzo1 causes fragmentation of the mitochondrial network. To avoid selecting against petite cells, we used synthetic growth medium containing amino acids and 1 % glucose (SC-glucose), where cells do not rely on respiration. Mitochondrial morphology was visualized using mCardinal targeted by preSu9, a well-established marker for mitochondria in yeast (Di Bartolomeo et al., 2020; Vowinckel et al., 2015). To quantify alterations of mitochondrial morphology, we segmented mitochondria in 3D and determined their solidity, a measure to describe the compactness of an object (Vőfély et al., 2019). Solidity is high for compact objects like fragmented or clumped mitochondria, while solidity is lower for wild-type like more tubular mitochondria. To determine the timing of mitochondrial morphology changes precisely, we performed confocal microscopy with a frame rate of 30 seconds. Alterations in mitochondrial morphology are detectable within 20 minutes after Fzo1 depletion and fragmentation is almost complete 60 minutes after Fzo1 depletion (Fig 1B and 1C and Video V1). In control cells, mitochondrial morphology is maintained throughout the imaged time course (Fig S1A). This observation is in line with studies showing that mitochondrial fusion and fission events occur every few minutes (Jakobs et al., 2003; Nunnari et al., 1997; Wisniewski et al., 2024). Disturbing this equilibrium by inhibition of one of the processes consequently results in a fast change of mitochondrial morphology.

We further confirmed that mitochondrial morphology is unaffected by TIR expression or the addition of the inducers (Fig S1B and S1C). Subsequently, we therefore only compared cells bearing TIR and FLAG-AID-Fzo1, which are either depleted or untreated (in the following termed Control).

After showing that mitochondrial morphology changes quickly upon Fzo1 loss, we sought to determine how long it takes wildtype (WT)-like cells to reach the full phenotype of Δ*fzo1* cells, which exhibit a growth defect on glucose media of ∼40 % compared to Control cells, consistent with previous reports (Shirozu et al., 2016). We found that loss of the Fzo1 protein results in a mildly decreased growth rate already within the first hours after onset of depletion. The decrease continues gradually reaching a growth rate comparable to Δ*fzo1* cells between 18-21 h, which corresponds to 10-12 doublings (Fig 1D, see Fig S1D for additional controls). Thus, while fragmentation of mitochondria, the primary phenotype of Fzo1 loss, is achieved within less than an hour, full reduction in growth is reached only after 21 hours.

### Loss of mtDNA and the respiratory chain occur together with alterations of mitochondrial ultrastructure

To understand the relationship between growth reduction and mtDNA loss after Fzo1 depletion, we determined mtDNA levels relative to nuclear DNA using quantitative DNA-PCR (qPCR). The depletion of Fzo1 leads to a severe loss of mtDNA within a few generations (Fig 2A), following an exponential decline with a half-life of approximately 4.5 hours (Fig S2G). We compared the levels of the different coding regions of the mitochondrial genome that had been described to be differentially lost upon ROS stress (Stenberg et al., 2022). However, we could not see differences between them (Fig S2A), suggesting that there is no loss of mtDNA segments prior to complete loss of mtDNA.

**Figure 2:**
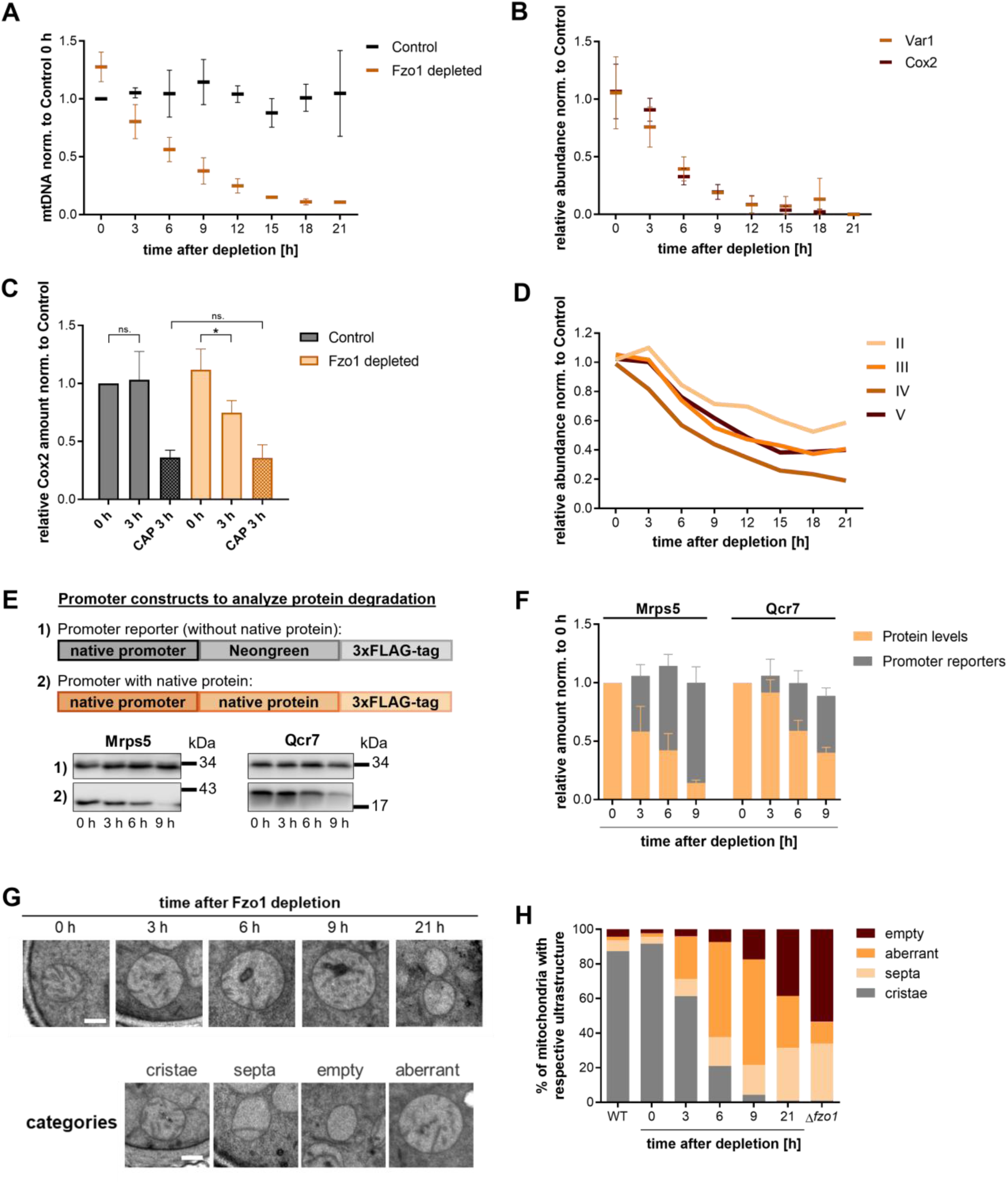
mtDNA is lost rapidly after Fzo1 depletion. A) DNA-qPCR of Fzo1 depleted and Control cells. mtDNA per nuclear DNA values were normalized to 0 h Control samples. B) Proteomics measurement of mtDNA encoded Var1 and Cox2. Peptide counts were normalized to 0 h Control samples. C) Quantification of Western Blot analysis of Cox2. Control and Fzo1 depleted cells were treated with 2 mg/mL chloramphenicol (CAP) to inhibit mitochondrial translation. Statistical significance was determined using an unpaired two tailed t-test. * indicate p < 0.05, n.s. = not significant. D) Proteomics measurement of nuclear encoded respiratory chain proteins. Lines represent the average of each complex. Abundances were normalized to 0 h Control cells. E) Western Blot analysis of 3xFLAG-tagged protein and promoter reporter protein for two nuclear encoded proteins required for mitochondrial respiration and translation. F) Quantification of Western Blots shown in E). G) Example images of electron microscopy, see Fig S2J for wildtype and Δ*fzo1* images. H) Quantification of electron microscopy images shown in G), scale bar = 200 nm. At least 150 mitochondria were analyzed for each time point. All measurements are mean values from three biological replicates. Where shown, error bars indicate SD.

We next wondered how rapidly mtDNA encoded proteins are lost. mtDNA in yeast encodes four subunits of the respiratory chain, three subunits of the ATP synthase (complex V) and one subunit of the mitochondrial ribosome. To analyze how fast the levels of these and other mitochondrial proteins decline, we performed mass-spectrometry based proteomics of the same time course as described above. Numbers of detected peptides and protein groups were comparable for all time points (Fig S2B). We first confirmed that upon start of the Fzo1 depletion (0 h), cells have a comparable proteome as Control cells in which TIR is not expressed, and that cells after 21 h of Fzo1 depletion have a proteome that is comparable to Δ*fzo1* cells (Fig S2C).

Of the mtDNA encoded proteins, Cox2 and Var1 were measured with high reproducibility across time points. Interestingly, the rate with which the abundance of Cox2 and Var1 proteins decreases was similar to the rate of mtDNA loss (Fig 2B), which we also confirmed by Western Blot analysis of Cox2 (Fig S2D). Since there was no delay between the decline of mtDNA and mtDNA encoded proteins, we investigated if mtDNA encoded proteins are actively degraded. We inhibited mitochondrial translation using chloramphenicol (CAP) in Control and Fzo1 depleted cells, in a Δ*pdr5* deletion strain which is defective in CAP export from the cell, thereby allowing efficient translational inhibition (Leonard et al., 1994; Roussou et al., 2024). As in previous experiments, we detected a decrease in Cox2 abundance 3 h after Fzo1 depletion (Fig 2C). Inhibition of mitochondrial translation resulted in a strong decrease of Cox2 in Control cells which correlates with dilution through growth. Fzo1 depleted cells treated with CAP showed a similar decrease in Cox2 abundance (Fig 2C, Fig S2E). This indicates that active degradation of mtDNA encoded proteins does not play a major role in the loss of mtDNA encoded proteins, but instead their synthesis is strongly reduced. In line with the reduction of synthesis of mitochondrial encoded proteins, we also observe a strong reduction of mitochondrial ribosomal proteins, while cytosolic ribosomal proteins were not reduced (Fig S2F).

The levels of nuclear-encoded respiratory chain proteins are known to be coupled to the presence of mtDNA encoded respiratory chain proteins (Dagsgaard et al., 2001; Vowinckel et al., 2021). In line with this, most nuclear encoded subunits of the respiratory chain complexes also decrease after loss of Fzo1 (Fig 2D), although with a slight delay compared to mtDNA encoded subunits (Fig 2B). Our proteomics analysis confirms that most of the nuclear encoded respiratory chain proteins are strongly reduced in Δ*fzo1* cells (Fig S2H).

We next sought to investigate whether the reduction in nuclear-encoded respiratory chain and mitochondrial ribosomal proteins is due to decreased synthesis through retrograde signaling or whether they are synthesized at a constant rate and subsequently degraded. We first confirmed by Western blot that two representative endogenously Flag-tagged proteins, Mrps5 and Qcr7, decrease. We then constructed reporters consisting of 800 bp of the respective promoter regions of these proteins driving the expression of Neongreen-FLAG (Fig 2E). While the abundance of the proteins (Fig 2E,F) are reduced by 60 – 80 % within the first 9 h after depletion, the abundance of the promoter-Neongreen constructs are stable (Fig 2E and 2F). This shows that contrary to mtDNA encoded proteins, nuclear encoded proteins involved in respiration are actively degraded.

The ATP synthase is composed of nuclear and mitochondria encoded subunits and is strongly reduced upon Fzo1 depletion (complex V, Fig 2D). The formation of ATP synthase dimers is required for normal cristae structure (Klecker & Westermann, 2021; Paumard et al., 2002). In line with this, we observe changes in mitochondrial ultrastructure after Fzo1 depletion (Fig 2G and 2H). In the first 3 h following depletion, most mitochondria still have cristae. However, 6 h and 9 h after Fzo1 depletion, most mitochondria have aberrant ultrastructure and 21 h after depletion most mitochondria resemble the Δ*fzo1* deletion phenotype, where mitochondria show a complete loss of normal cristae (Fig 2G and 2H, see Fig S2J for wild-type and Δ*fzo1* examples). During the first 9 h of depletion, many cells have a mixture of WT and deletion-like mitochondria (Fig S2K) suggesting that loss of functionality and likely mtDNA is not homogenous, not even within individual cells.

### Fzo1 depleted cells exhibit large variation in mitochondrial mass and functionality

Since respiratory chain proteins were strongly reduced, we next investigated if mitochondrial content in the cell is generally affected after Fzo1 depletion. Mitochondrial biosynthesis in WT cells is regulated with cell size, which leads to a roughly constant concentration of mitochondria (total volume of mitochondria per cell volume) in the population (Rafelski et al., 2012; Seel et al., 2023). 3-dimensional volume reconstruction of mitochondria may be biased by the different shapes of WT and fragmented mitochondria. Therefore, we instead used the total fluorescence signal intensity derived from mitochondrial imported pre-Su9-mCardinal and normalized it to the cell volume as a robust approximation of mitochondrial concentration (in the following termed mitochondrial concentration, see also Methods and Fig S3A, B for controls).

We show that the mean of the mitochondrial concentration of the population does not decrease within the first 9 h after Fzo1 depletion (Fig S3C). However, the single-cell distribution of mitochondrial concentration broadens substantially, and at the same time becomes less normally distributed. This results in a decrease of the median mitochondrial concentration approximately 1 h after depletion (Fig 3A), leading to a large fraction of the population with an extremely low mitochondrial concentration as depicted by the 25^th^ percentile of the population (Fig 3A). This becomes even more pronounced if Fzo1 is depleted for a prolonged period of time (Fig S3D).

**Figure 3:**
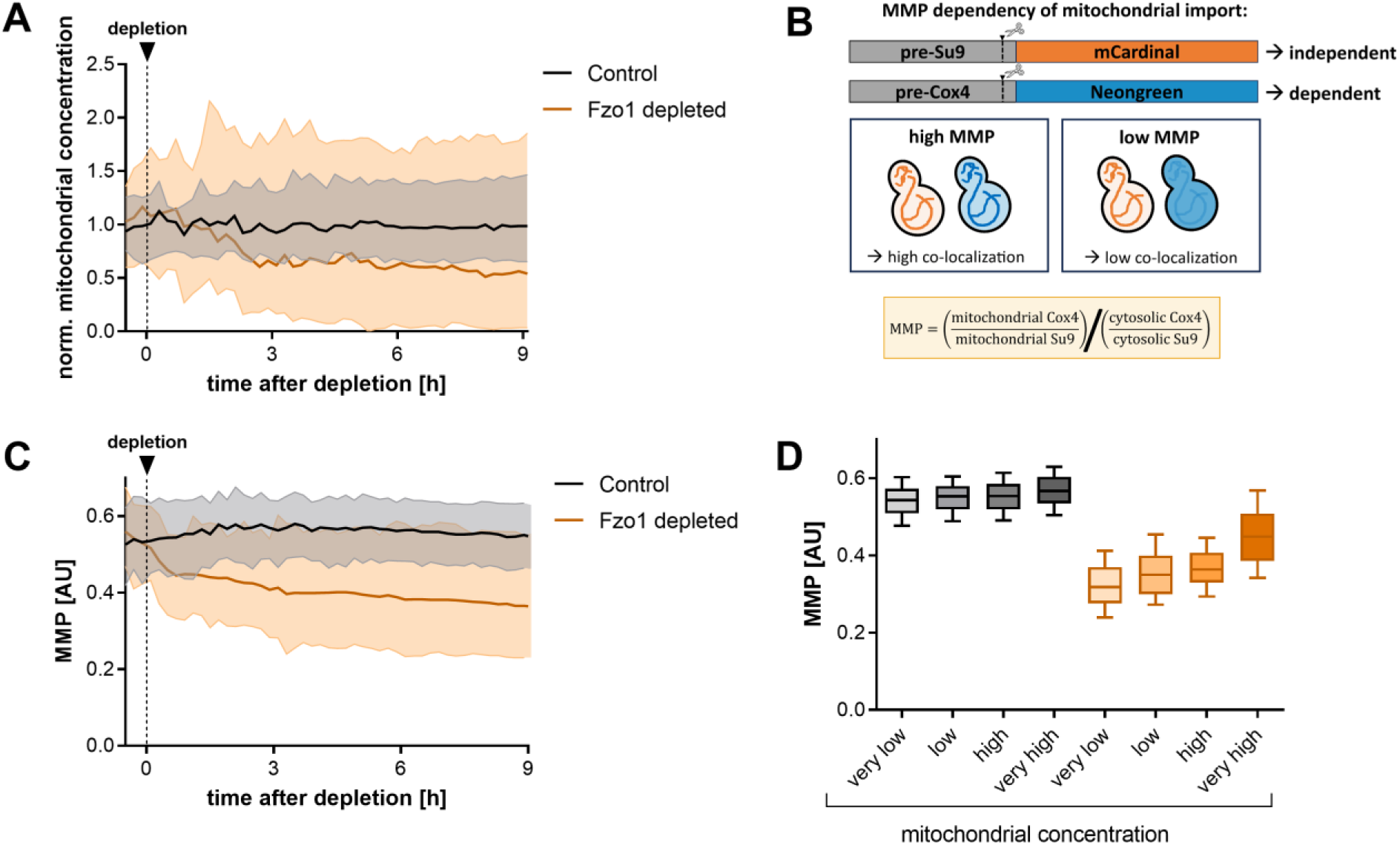
Fzo1 depleted cells exhibit large variation in mitochondrial concentration and functionality. A) Normalized mitochondrial concentration (total mitochondrial signal intensity per cell volume). Depletion is induced at t = 0. Median with 25^th^ and 75^th^ percentiles from two biological replicates with a total of 1785 Control and 2075 Fzo1 depleted cells is shown. B) Schematic of the reporter for estimating mitochondrial membrane potential (MMP). Mitochondrial import of pre-Su9 is mostly independent of the MMP while import of pre-Cox4 is dependent on the MMP. C) Estimation of the MMP after Fzo1 depletion. Average with 5^th^ and 95^th^ percentiles from two biological replicates with a total of 1723 Control and 1595 Fzo1 depleted cells is shown. D) MMP of cells with different mitochondrial concentrations of all G1 cells 0 – 9 h after Fzo1 depletion. Categories were determined by the quartiles of the mitochondrial concentrations of Control cells. Medians with 10^th^ and 90^th^ percentiles are shown. Control cells: 407 cells per category, Fzo1 depleted cells: n (very low) = 821, n (low) = 184, n (high) = 183, n (very high) = 581.

Since WT cells maintain relatively constant mitochondrial concentrations (Rafelski et al., 2012; Seel et al., 2023), we investigated if this large variability of mitochondrial concentrations after Fzo1 depletion affects mitochondrial function, such as the mitochondrial membrane potential (MMP). To estimate MMP, we used the Mitoloc system (Vowinckel et al., 2015), which consists of two fluorescent reporters: one containing the Su9 presequence whose import into mitochondria is largely independent of the MMP, and one containing the Cox4 presequence whose import is strongly dependent on the MMP (Fig 3B). The ratio of the two reporters in the mitochondria thus allows an approximation of the MMP. We improved the signal-to-noise of the original construct by integrating the reporter into the genome and expressing both fluorophores from the same strong constitutive promoter. Before Fzo1 depletion, the MMP is comparable to Control cells. Following the MMP over time revealed that the MMP shows an immediate drop after Fzo1 depletion, simultaneously with the mitochondrial network fragmentation. The MMP then decreases further within the first 9 h of depletion and reaches a deletion-like level between 21 – 24 h (Fig S3E,F), while Control cells maintain a stable MMP. Interestingly, the MMP of Fzo1 depleted cells also shows an increased variability between individual cells, indicating a large variation in mitochondrial function (Fig 3C) that is in line with the observed heterogeneity in ultrastructure.

Since we detected a broadened distribution for both mitochondrial concentration and MMP, we checked whether there is a correlation between mitochondrial concentration and MMP. We grouped G1 Control cells based on their mitochondrial concentration into quartiles. Control cells show very similar MMP values independent of their mitochondrial concentration. In contrast, using the same categories determined from Control cells, we find that Fzo1 depleted cells show large differences in the MMP depending on their mitochondrial concentration. Cells with low mitochondrial concentrations exhibit a much lower MMP than those with high mitochondrial concentrations (Fig 3D). Mitochondrial health is known to be important for cellular health. In line with this, we found that cells with low mitochondrial concentrations exhibit extended division times compared to those with higher mitochondrial concentrations (Fig S3G). This is most strongly pronounced during later times of depletion (8-18 h).

### Fzo1 depleted cells exhibit mitochondrial inheritance defects

Having shown that Fzo1 depletion leads to a large variability in mitochondrial concentration (total mitochondrial signal intensity per cell volume) which correlates with mitochondrial MMP, we investigated which cells obtain a low mitochondrial concentration. Since it was known that Δ*fzo1* deletion cells exhibit delayed mitochondrial inheritance (Böckler et al., 2017), we analyzed if buds inherit less mitochondria in Fzo1 depleted cells. We determined the size at which buds first receive mitochondria in Δ*fzo1* and Fzo1 depleted cells. Δ*fzo1* and Fzo1-depleted buds receive mitochondria on average at a larger size and at later timepoints than Control cells (Fig 4A, B, Fig S4A), confirming observations from Böckler et al.

**Figure 4:**
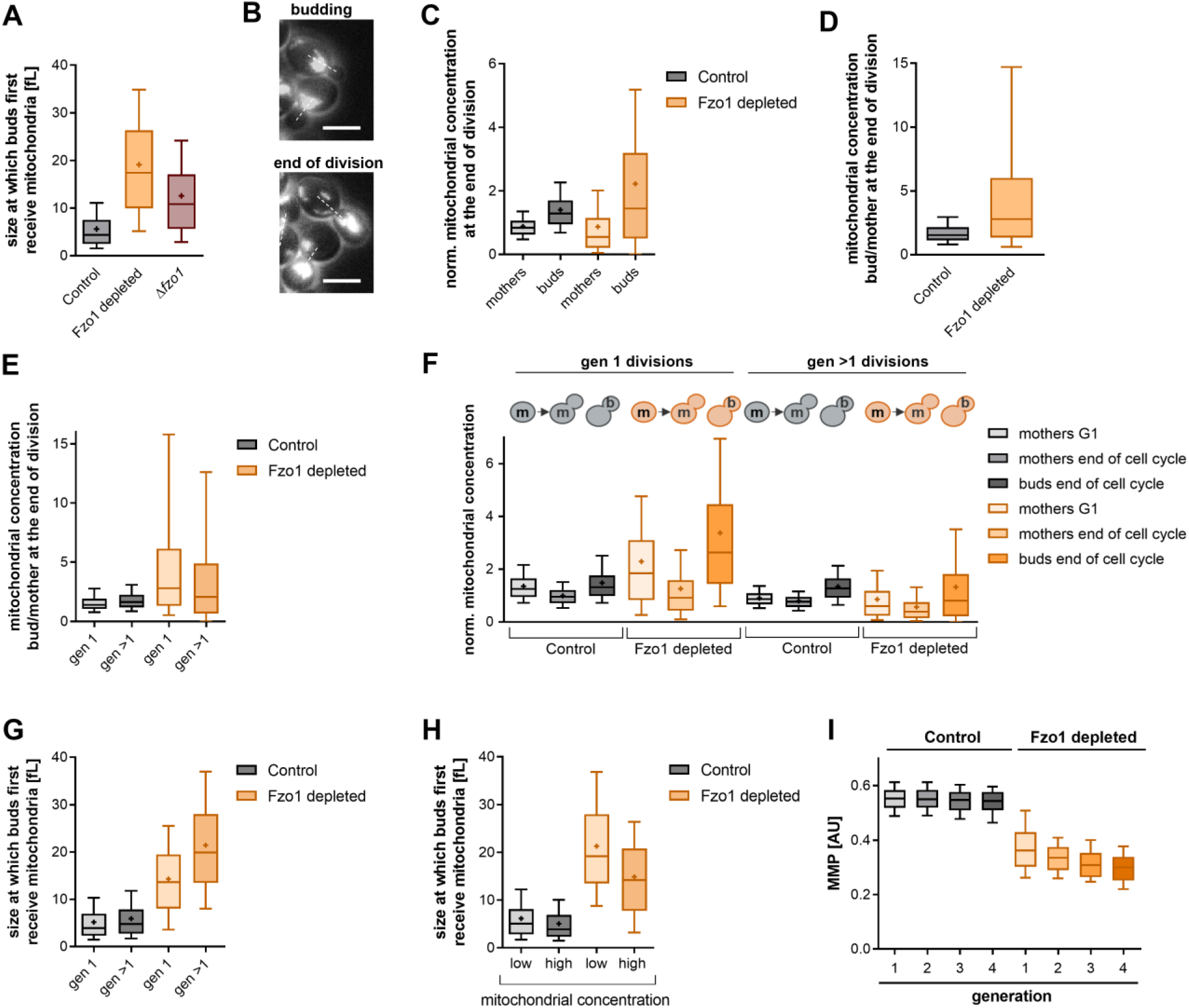
Fzo1 depleted cells exhibit mitochondrial inheritance defects. A) Cells size at which buds first receive mitochondria. Images were taken every 8 min and the time preSu9-mCardinal was detected the first time in the bud was determined. Control = 1249, Fzo1 depleted = 1275, Δ*fzo1* = 1237. B) Example images of mitochondria after Fzo1 depletion during budding and end of the cell cycle. Maximum z-projections are shown. C) Normalized mitochondrial concentration (total mitochondrial signal intensity per cell volume) at the last recorded timepoint before division. n (Control) = 1250, n (Fzo1 depleted) = 1409. D) Ratio of the mitochondrial concentration of buds to mothers at the end of the cell cycle of cells shown in C). E) As D) for gen 1 and gen >1. Control: n (gen 1) = 544, n (gen >1) = 706, Fzo1 depleted: n (gen 1) = 586, n (gen >1) = 735. F) Mitochondrial concentrations of mothers and buds through the cell cycle of gen 1 and gen >1 divisions. Number of cells as in E). G) Size at which buds of gen 1 or gen >1 mothers first receive mitochondria. Number of cells as in E). H) Size at which buds of mothers with low or high mitochondrial concentrations first receive mitochondria. The median mitochondrial concentration of Control cells at G1 was used to determine low and high categories. Control: n (low) = 625, n (high) = 624, Fzo1 depleted: n (low) = 638, n (high) = 637. I) Estimation of MMP as described in Fig 3 of generation 1-4 mothers at the beginning of the cell cycle. Control: n (gen 1) = 1478, n (gen 2) = 383, n (gen 3) = 215, n (gen 4) = 114, Fzo1 depleted: n (gen 1) = 1550, n (gen 2) = 406, n (gen 3) = 220, n (gen 4) = 102. See Fig S4C for mitochondrial concentrations.

However, when following mitochondria to the end of the cell cycle, a different picture emerged. Immediately before division, a much higher mitochondrial concentration was detected in buds compared to mothers (Fig 4B and 4C). We determined the ratio of mitochondrial concentration of each bud in relation to its mother at the end of the cell cycle. Fzo1 depleted cells show a 60 % higher median ratio and a higher variability of this ratio compared to Control cells. (Fig 4D). This was also observed when Fzo1 was depleted for prolonged periods of time and in Δ*fzo1* cells (Fig S4B). Thus, even though buds of Fzo1 depleted cells receive mitochondria later than buds of Control cells, they often receive a much higher mitochondria concentration at the end of the cycle.

Given that at division buds typically have a higher concentration of mitochondria than their mothers, we wondered how this would affect mitochondria during the following cell cycles. After division, the bud becomes a ‘first-generation mother’, while its mother continues with its next cell cycle. In first-generation mothers, Fzo1 depleted cells start off with a higher mitochondria concentration than first-generation Control cells (Fig 4F). However, these first-generation mothers then mostly transfer more mitochondria to their bud than they retain, resulting in a higher mitochondrial concentration ratio between the bud and mother at the end of the cell cycle (Fig 4E) and a strong reduction of the mitochondrial concentration of the mother (Fig 4F) compared to Control cells.

In higher-generation divisions, where the mother typically already starts off with a lower mitochondrial concentration, the median ratio of the mitochondrial concentration between bud and mother is lower than in first generation divisions but still higher than in Control cells. However, in these higher generation divisions the distribution of the ratios shifts, so that in some divisions, the buds obtain a very low mitochondrial concentration (Fig 4E and F). We next asked why buds of higher generation divisions sometimes obtain very little mitochondria. This may be caused by the fact that buds of higher generation mothers, which often have a low mitochondrial concentration compared to first generation mothers, obtain mitochondria at a larger size (Fig 4G,H) and at a later time (Fig S4E).

In line with a higher mitochondrial concentration in first generation cells compared to higher generation cells (Fig S4C), we measured a higher MMP in first than in higher generation cells (Fig 4I). Even at the same mitochondrial concentration, first generation cells have a higher MMP. While this effect can be clearly seen in Fzo1 depleted cells, Control cells have very similar MMP values throughout generations (Fig S4D). This indicates that not only a large amount, but also high-quality mitochondria are preferentially transferred into buds which will then become first generation mothers, and that mothers are left with a low amount of low-quality mitochondria.

Taken together, our results show that despite the delayed inheritance of mitochondria, buds obtain a large proportion of mitochondria at division. This causes a strongly unequal distribution of mitochondrial mass within the population, which in turn affects MMP and likely functionality.

### mtDNA encoded proteins are also segregated unequally during division

Having shown that mitochondrial mass is strongly unequally distributed through cell division with effects on mitochondrial functionality, we sought to investigate how this affects mtDNA encoded proteins. We therefore integrated the system to deplete Fzo1 in a previously established strain in which Atp6, a mtDNA encoded subunit of the ATP synthase, is tagged with mNeongreen in the mitochondrial genome (Jakubke, 2021). This was shown previously to be a good proxy for functional mtDNA (Roussou et al., 2024). We determined the Atp6 amount (total intensity) per cell volume and normalized this to the values immediately before Fzo1 depletion. Unlike the mean mitochondrial concentration which stays constant within the first 9 h, the mean Atp6 concentration already starts to decrease around 3 h after depletion and reaches about 20 % of initial levels at 9 h (Fig S5A). We ruled out a major role of mitophagy (Fig S5C and S5D) to this rapid drop. Rapid Atp6 reduction is in agreement with the changes of the other mtDNA encoded proteins measured by Western blot and proteomics (Fig 2B, S2D). 3 h after Fzo1 depletion, the median Atp6 concentration has decreased to 50 % compared to control cells and further decreases within the first 9 h (Fig 5A and 5B).

**Figure 5:**
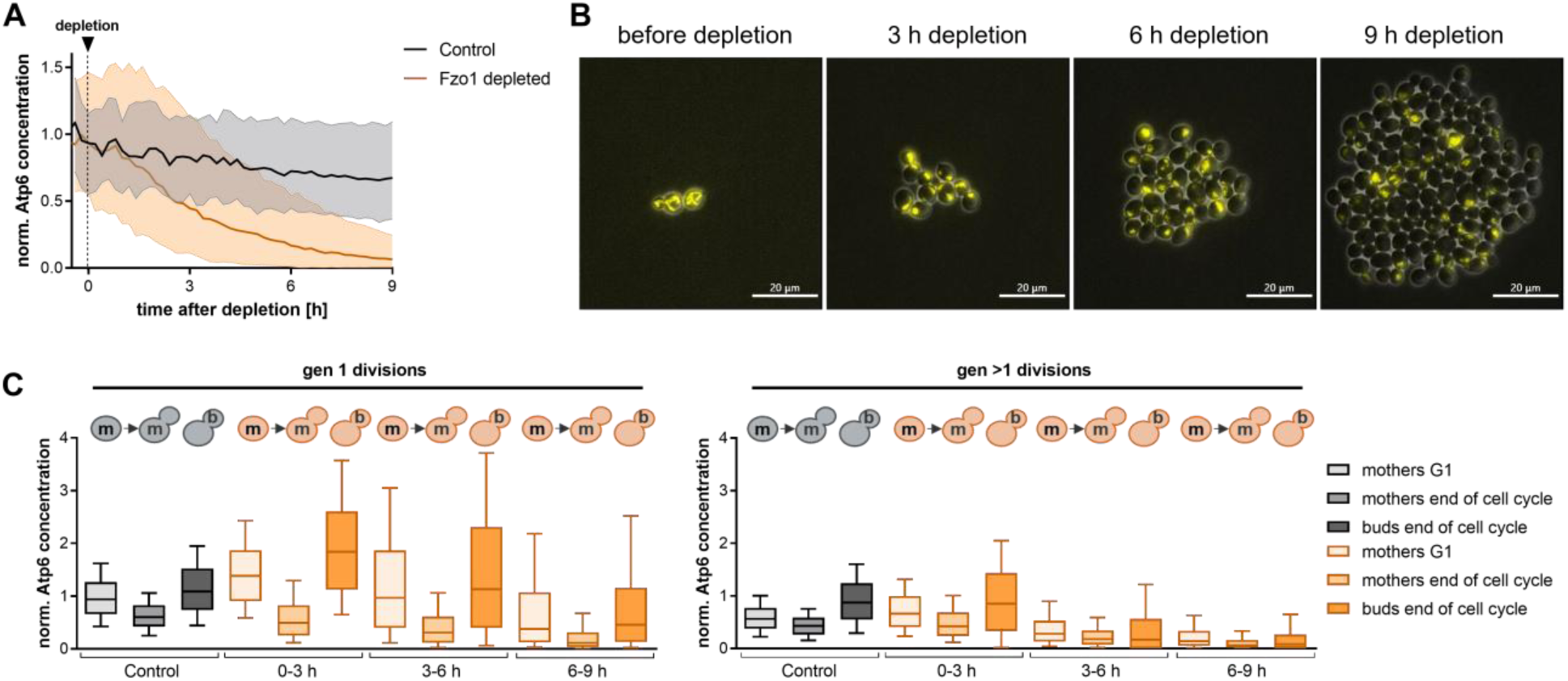
mtDNA encoded proteins are segregated unequally during division. A) Atp6-mNeongreen concentration (total mitochondrial mNeongreen intensity per cell volume) after Fzo1 depletion. Median with 25^th^ and 75^th^ percentiles is shown. Number of cells at 9 h: Control = 3818, Fzo1 depleted = 3686. B) Example images of Atp6-mNeongreen. Scale bar = 20 µm. C) Atp6 concentration of mothers at the beginning of the cell cycle (G1), and mothers and buds at the end of the cell cycle. n (gen 1): Control = 1169, 0-3 h = 592, 3-6 h = 2384, 6-9 h = 844, n (gen >1): Control = 1486, 0-3 h = 865, 3-6 h = 3236, 6-9 h = 1092.

Since we observed a disbalance in the distribution of mitochondrial concentrations between buds and their mothers, we examined if Atp6 concentration is also unequally distributed through division. We analyzed the Atp6 concentration of mothers and their buds at the end of the cell cycle. In first-generation divisions, the Atp6 concentration drops drastically in mother cells while their buds obtain as much as the mothers had at the beginning of the cell cycle (Fig 5C, left). In higher generation divisions, the Atp6 concentration of the mother also drops through cell division. While the median Atp6 concentration at the end of the cell cycle is also slightly higher in buds than in mothers, many buds do not receive any Atp6. Often, the Atp6 content present at the beginning of the cell cycle is distributed between mother and bud, resulting in a much lower concentration in both the mother and the bud at the end of division (Fig 5C, right, Fig S5B). This implies that higher generation cells do not synthesize a sufficient amount of Atp6 during the cell cycle to supply both mothers and buds with an appropriate concentration of Atp6. This is likely also the case for the other mtDNA-encoded proteins.

### Compensatory synthesis of mtDNA-encoded protein fails after Fzo1 depletion

To determine if first or higher generation mothers exhibit differences in the amount of Atp6 they are able to synthesize, we calculated the change of Atp6 over a full cell division cycle. We subtracted the amount of Atp6 from the mother at the beginning of the cell cycle from summed Atp6 amount of the mother and daughter at the beginning of the next G1. First generation cells are able to synthesize Atp6 after Fzo1 depletion, but exhibit a continuous reduction in the amount of Atp6 produced within one cell cycle within the first 9 h after Fzo1 depletion. Higher generation cells produce almost no Atp6 already 5 h after Fzo1 depletion (Fig 6A, Video V2), indicating that they reach a deletion-like state faster than first generation cells.

**Figure 6:**
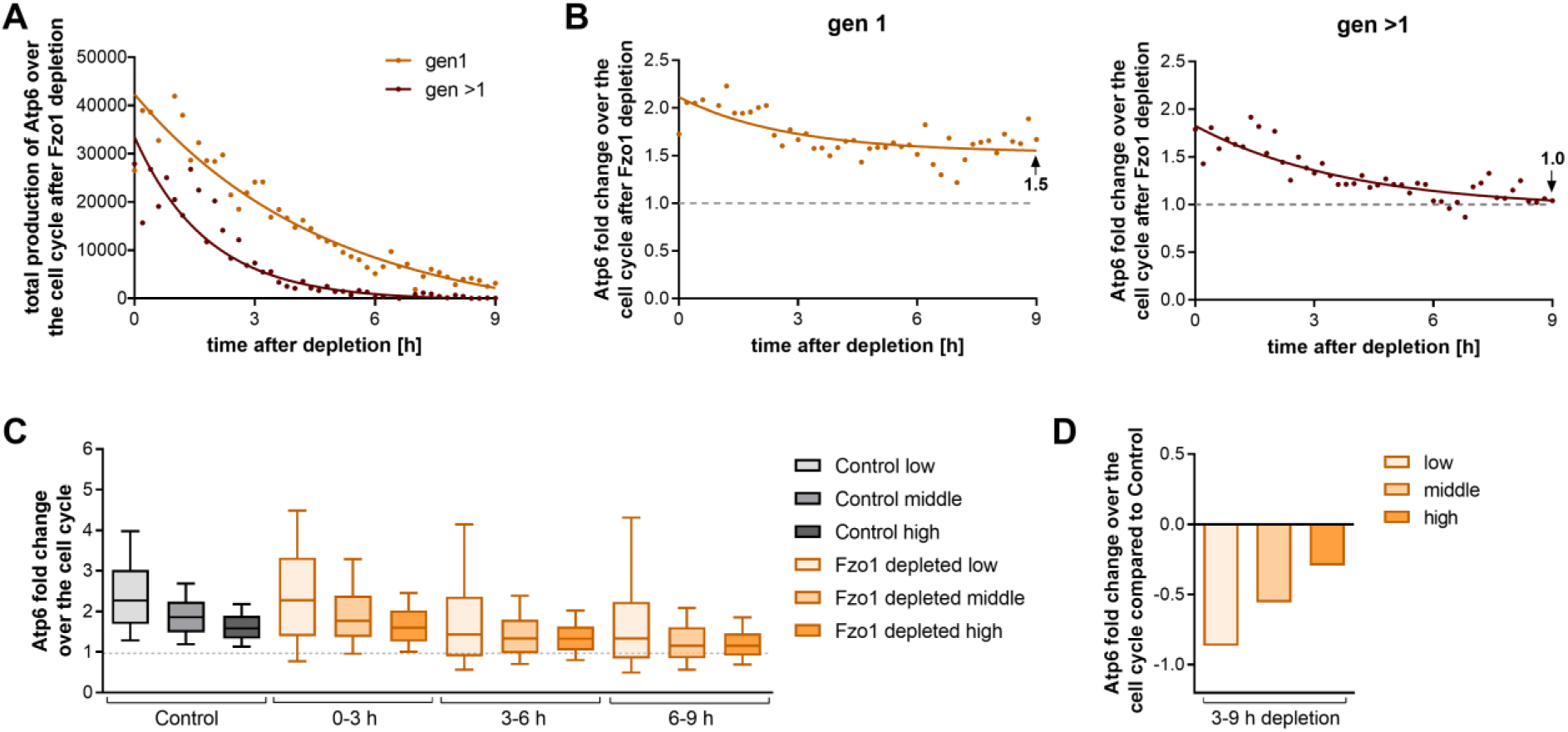
synthesis of mtDNA encoded proteins is reduced. A) Total change of Atp6-mNeongreen (arbitrary fluorescence intensity units) through the cell cycle of first and higher generation divisions. 3820 divisions for gen 1 and 5193 divisions for gen >1 divisions were analyzed. The median of each timepoint is shown. B) Fold change of Atp6-mNeongreen through the cell cycle of first and higher generation divisions as shown in A). C) Fold change of Atp6 over the cell cycle of divisions of cells with low, middle, and high Atp6 content of cells shown in A) & B). Control (whole timeframe): each category: n = 831. Fzo1 depleted 0 – 3 h: n (low) = 276, n (middle) = 212, n (high) = 964, Fzo1 depleted 3 – 6 h: n (low) = 3006, n (middle) = 774, n (high) = 1672, Fzo1 depleted 6 – 9 h: n (low) = 1375, n (middle) = 172, n (high) = 296. D) Difference of the median relative synthesis of Atp6 through cell divisions of Fzo1 depleted cells compared to Control cells of 3 – 9 h after Fzo1 depletion from data shown in C).

We next determined the fold change of Atp6 over the cell cycle over time. If a cell produces exactly enough new mtDNA to maintain mtDNA concentrations over the cell cycle, then the expected fold-change value would be close to two (a little more in first-generation mothers since the total volume of the mother plus bud is more than twice the volume of a newborn cell). Before depletion, the fold change of Atp6 is close to this expected value but decreases gradually over time after Fzo1 depletion reaching 1.3, averaged across generations at 9 h (Fig S6A), while the fold change of imported preSu9-mCardinal over the cell cycle stays almost constant (Fig S6B). However, first generation cells at this time are still able to produce Atp6 with a fold change of 1.5 while higher generations are at 1.0, indicating that Atp6 is not synthesized and only diluted in these divisions (Fig 6B). This again indicates that higher generation cells likely become petite earlier than first generation cells.

Since we observed a strongly unequal distribution of Atp6 within the population (Fig 5A,B) we wanted to investigate if the amount of Atp6 produced in a given cell is dependent on the initial amount of Atp6 at the beginning of its cell cycle. We therefore determined categories by dividing the Control population at the beginning of G1 into three equally large groups: low, middle and high Atp6 amount. In Control cells, the cells with low Atp6 amount have a higher relative production of Atp6 over the cell cycle, while those with high Atp6 amount have a much lower relative synthesis (Fig 6C and Fig S6D). This indicates that in WT cells a feedback mechanism exists, ensuring homeostasis through reduced or increased synthesis, similar to the mechanism observed in fission yeast (Jajoo et al., 2016). Fzo1 depleted cells lose this compensatory mechanism. Between 3-9 h after depletion a reduction in the Atp6 fold change over the cell cycle compared to Control cells can be detected for all three categories (Fig 6D). However, the difference compared to Control cells is most strongly pronounced in cells with low Atp6 content. While Control cells with low Atp6 content exhibit a ∼2.5 fold change of Atp6 over the cell cycle to compensate for the low starting amount, Fzo1 depleted cells only exhibit a 1.4 fold change between 3-9 h after depletion. This suggests that the compensatory mechanism that ensures maintenance of mtDNA fails in Fzo1 depleted cells. Through the shift of the fraction of the population with extremely low and high Atp6 amount, the unequal distribution further enhances a reduction in the Atp6 fold change (Fig S6D-F). In conclusion, these results suggest that the combined effect of the unequal distribution and reduction in synthesis results in a reduction of Atp6 over time.

### Mathematical modeling shows that unequal distribution of mitochondria in combination with the reduction of synthesis leads to a fast and complete establishment of the Δ*fzo1* phenotype

Our experiments showed that depletion of Fzo1 affects the inheritance of mitochondria to the bud at cell division. At the same time, it leads to decreased relative synthesis rate of mtDNA encoded Atp6 during the cell cycle, in particular for cells with low starting concentrations of Atp6. Since we consider Atp6 to be a good proxy for the amount of functional mtDNA, this decreased rate is likely due to a decreased synthesis of mtDNA. To dissect how these two effects contribute to loss of mtDNA, we built a simple model simulating mtDNA inheritance and synthesis over many cell cycles.

For each simulation, we initiated a population of 10 000 cells, each starting with 16 nucleoids, the structures that contain mtDNA (Roussou et al., 2024), and each about to divide. We then assumed that each cell would distribute all its nucleoids between mother cell and bud with a ratio randomly drawn from log-normal distribution, rounded to integers. The parameters for the log-normal distribution were experimentally determined for both Control cells (during the entire time period) and Fzo1-depleted cells (3-9 h after onset of depletion) (Fig S7A). After cell division, the number of total cells was randomly reduced by 50% to again obtain 10 000. Next, we simulated mtDNA synthesis during the cell cycle, where the amount of synthesis depends on the initial nucleoid number, which was sorted in five groups from very low to very high (Fig S7B, see Methods and Figure Caption for further details). The values for this relative synthesis rate were determined from experiments and were different for Control and Fzo1-depleted cells (Fig S7B). Continuing with simulation of the next cell division, this procedure was then repeated for a total of 100 cell cycles.

We show that the segregation of mtDNA between mother and bud as approximated by Atp6 measurements, together with the five categories of mtDNA/Atp6 fold change over the cell cycle are sufficient to recapitulate mtDNA homeostasis in WT-like cells, as the simulated levels for Control cells are stable over time (Fig 7). Contrary to this, Fzo1 depletion-like segregation of nucleoids between mother and bud combined with Control cell Atp6 synthesis, result in a continuous but slow decrease of nucleoid number over time. On the other hand, WT-like segregation combined with Fzo1 depletion-like synthesis leads to a rapid decrease in nucleoid numbers within the first divisions, followed by a very slow decrease during the following divisions. Only the combined effect of both, the unequal distribution and reduced synthesis leads to loss of nucleoids with dynamics that match those we observed experimentally. Since for the simulation we used the Atp6 synthesis data obtained between 3-9 h after Fzo1 depletion and the synthesis likely drops further during later times of depletion, we expect the decrease of nucleoids to be even faster than predicted by our simulation.

**Figure 7:**
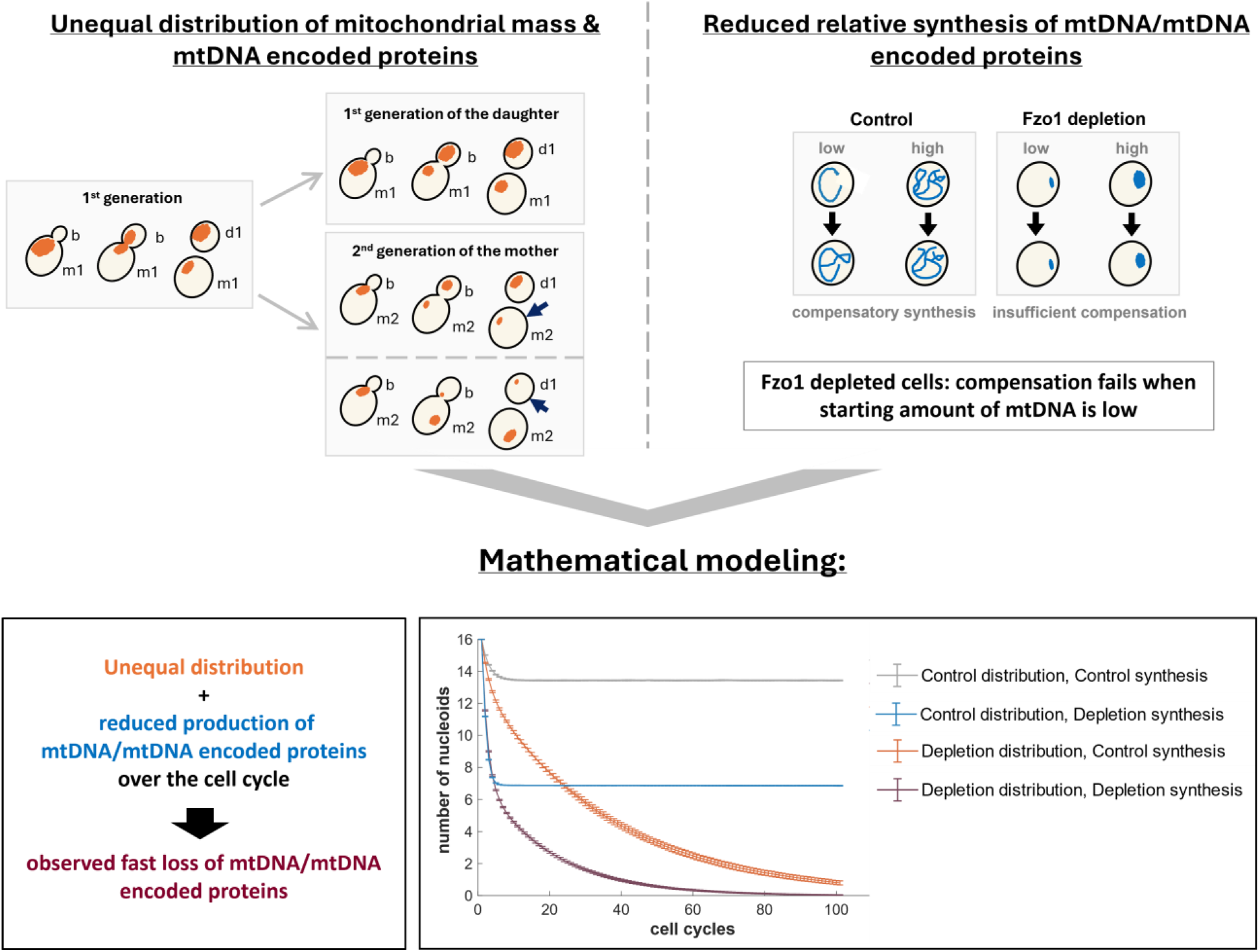
The combination of unequal segregation and reduced synthesis is necessary and sufficient for the Δ*fzo1* phenotype development. Schematic depiction of observed phenotypes: left: unequal distribution, right: reduced synthesis of mtDNA/mtDNA encoded proteins. Below: mtDNA level modeled with Control or Fzo1 distribution and synthesis of mtDNA encoded Atp6. See Fig S7 and methods for model assumptions and parameters.

Taken together, our mathematical model shows that both factors, the unequal distribution of mtDNA and a reduction in relative synthesis, are required and also sufficient to explain the development of the Fzo1 phenotype.

## Discussion

Here, we unravel how mtDNA is lost in cells lacking an active mitochondrial fusion machinery. We used the AID-system to rapidly deplete the mitochondrial outer membrane mitofusin Fzo1 in combination with microfluidics-based, quantitative single-cell time-lapse imaging of tens of thousands of cells across their cell cycles. With these tools, we could resolve the timing of the individual defects that lead to loss of mtDNA: altered mitochondrial morphology, reduction of the mitochondrial membrane potential (MMP), and reduced synthesis of mtDNA and mtDNA-encoded proteins.

In line with a previous study (Hermann et al., 1998), we show that fragmentation of the mitochondrial network is the primary phenotype of Fzo1 depletion and occurs in less than one hour. Simultaneously with mitochondrial fragmentation, we observed an initial drop in mitochondrial membrane potential (MMP), which decreased further until 21 h of Fzo1 depletion. Approximately at the time MMP reaches an Δ*fzo1* deletion-like state, also deletion-like growth is reached.

Surprisingly, most of the mtDNA (Fig 2A) and mitochondrially encoded proteins (Fig 2B) were lost within the first 9 h of Fzo1 depletion. At a comparable timescale to the loss of mtDNA and the proteins encoded by it, we observed a reduction of nuclear-encoded respiratory chain proteins. This results in a strong decrease of ATP synthase, which is required for normal mitochondrial ultrastructure. Consequently, we detected a stepwise change in mitochondrial ultrastructure from mostly cristae to a mixture of cristae and aberrant mitochondria to an ultrastructure that resembles petite cells. As observed for other parameters, mitochondrial ultrastructure resembles that of Δ*fzo1* cells after 21 h of depletion.

After 21 hours, the Fzo1 depletion shows changes comparable to Δ*fzo1* cells in the proteome, such as downregulation of respiratory chain proteins and upregulation of glycolytic enzymes (Fig S2I). These larger-scale changes in the proteome only start approximately 9 h after Fzo1 depletion, when most of the mtDNA has already been lost (Fig S2I). These proteome changes are most likely a generic response to the metabolic deficiencies caused by absence of mtDNA, rather than specifically to Fzo1 depletion, as petite cells obtained by ethidium bromide treatment showed comparable proteome changes (Vowinckel et al., 2021). Supporting this, it was postulated that mitochondria-to-nucleus communication is mostly mediated by the cytosolic signalling machinery as a response to metabolic changes (Knorre et al., 2016). In line with an absence of signalling to the nucleus, we detected no changes in the promoter activity of nuclear genes encoding mitochondrial proteins involved in respiration within the first 9 h of depletion, even though the protein levels strongly declined.

While the growth limitations and metabolic stress reactions of the final phenotype of petites are similar regardless of how petites emerged, the mechanisms leading to the loss of mtDNA are likely to differ. The loss of mtDNA and the proteins encoded by it may be expected to be caused by mitophagy of low functioning mitochondria, but we could exclude a major role of mitophagy (Fig S5C and S5D). Also, mtDNA integrity does not seem to be affected since different loci on the mtDNA decreased with a similar rate, as also seen in mouse embryonic fibroblasts with a mitofusin deletion (Silva Ramos et al., 2019).

By analyzing the dynamics of mitochondria and mtDNA-encoded proteins on a single cell level using live-cell time-lapse imaging, we were able to identify the underlying cause of mtDNA loss: the combination of unequal inheritance of mitochondria and decreased synthesis of mtDNA. Despite the mitochondrial inheritance delay of Fzo1-depleted cells as described for Δ*fzo1* cells (Böckler et al., 2017), at the end of the cell cycle buds obtain a high concentration (total mitochondrial signal intensity per cell volume) of mitochondria, whereas mothers often only retain a low concentration compared to their buds. While the overall mitochondrial mass within the population is maintained within the first 9 h of depletion, this unequal distribution results in a progressive decline of mitochondrial health within the population. This is in line with a recent preprint (Ray et al., 2025) showing that cells which inherit low concentrations of mitochondria are more likely to become petite. Additionally, this unequal distribution also impacts mtDNA maintenance. Control cells maintain stable mean mtDNA levels across the population, even if starting levels in individual cells fluctuate (Fig 6C), suggesting a compensatory mechanism. In contrast, in Fzo1 depleted cells this compensatory mechanism fails. Cells with low amounts of mitochondrial encoded Atp6 show a strong reduction in synthesis rates compared to wild-type like cells with similar starting amounts, and can thus not recover if they do not inherit a sufficient amount of mtDNA and the proteins encoded by it. This combination of unbalanced inheritance and reduced synthesis is necessary and sufficient to explain the observed fast and complete loss of mtDNA in Fzo1-depleted cells in a simple mathematical model (Fig 7).

Why are Fzo1-depleted cells unable to compensate for the low levels of mitochondrial-encoded proteins through increased synthesis, unlike wild-type cells? We hypothesize that several factors contribute to this phenomenon: In WT cells mtDNA containing nucleoids are regularly spaced, but in Fzo1-depleted cells might cluster in some mitochondrial fragments. Clustering of nucleoids was shown to impact mtDNA replication in mouse embryonic fibroblasts lacking mitofusions through an imbalance in replisome components (Silva Ramos et al., 2019). Regular spacing of nucleoids was shown to avoid stochastic distribution effects during inheritance, becoming too strong during fission yeast cell division (Jajoo et al., 2016).

Mitochondrial fragmentation may not only lead to unequally distributed nucleoids, but also unequally distributed import complexes within individual mitochondrial fragments. Khan and colleagues (Khan et al., 2024) postulate that this unequal distribution of import complexes might contribute to the reduction of MMP in Fzo1 deletion cells. Supporting this hypothesis, we detected an immediate reduction of MMP concurrently with fragmentation of the mitochondrial network.

The MMP serves as the driving force for the import of nuclear-encoded mitochondrial proteins into mitochondria. A reduction in MMP, therefore, leads to impaired mitochondrial import. Deficiency in the import of mitochondrial proteins eventually results in their degradation in the cytosol (Liu et al., 2021). Since nearly all mitochondrial proteins are nuclear-encoded, this disruption compromises mitochondrial functionality. Coordinated and efficient import is particularly important for components of the respiratory chain, as these proteins are highly hydrophobic and require the simultaneous provision of mtDNA-encoded subunits for the assembly of functional complexes. The stability of mtDNA-encoded proteins is affected by nuclear-encoded ones (Contamine & Picard, 2000). Therefore, impaired mitochondrial import of nuclear-encoded proteins through MMP reduction likely affects the stability of mtDNA-encoded proteins in Fzo1-depleted mitochondria. Complete loss of mtDNA leads to a significant reduction in the levels of nuclear-encoded respiratory chain proteins (Dagsgaard et al., 2001; Vowinckel et al., 2021). This highlights the complex interdependence between the stability and biogenesis of mtDNA and nuclear-encoded components, which likely drives a progressive decline in mitochondrial function if not carefully balanced. As a result, both mitochondrial membrane potential (MMP) and mitochondrial content are diminished, ultimately contributing to a decline in cellular health and a slowing of cell cycle progression (Chacko et al., 2025; Gorospe et al., 2023; Jajoo et al., 2016; Vowinckel et al., 2021).

Taken together, we show here that mtDNA loss in the absence of mitochondrial fusion is a consequence of complex, intertwined mitochondrial defects, with stochastic inheritance playing a key role. The use of single-cell time-lapse microscopy analysis across multiple generations proved essential to uncover the causality behind the emergence of these phenotypes. This approach allowed us to show how mitochondria with altered morphology and functionality are segregated during asymmetric division. Asymmetric division in metazoans allows distinct fates for their progeny, which is also reflected in mitochondrial inheritance (Hinge et al., 2020). In human stem-like cells, for example, younger mitochondria are preferentially retained in the stem cell to preserve its properties (Hinge et al., 2020; Katajisto et al., 2015). Similarly, undifferentiated lymphocytes preferentially inherit the youngest mitochondria, while senescent and differentiated lymphocytes receive the oldest mitochondria (Adams et al., 2016). By revealing how mitochondrial defects affect inheritance, our study enhances the understanding of mitochondrial segregation during asymmetric division with relevance to all types of asymmetric divisions where mitochondrial morphology or function is altered.

## Author Contributions

LD and JCE conceived and designed the study. LD performed most experiments and data analysis. FP provided code and supported data analysis. BL performed and analyzed proteomics experiments. RB performed and analyzed electron microscopy. KMS supported study design, data analysis and interpretation, and developed the mathematical model with input of LD. JCE, KMS, BM, and BW supervised the projects. LD wrote the first draft of the manuscript. JCE, KMS, BW edited the manuscript. All authors reviewed and approved the manuscript.

## Acknowledgments

We thank Masato Kanemaki, Christof Osman, Markus Ralser, Serge Pelet, Jan Skotheim, and Jörg Stelling labs for strains, plasmids, and reagents. We thank Luisa Hernández Götz, Helin Ezgi Çullu and Annika Duczmal for experimental support during their studies. We gratefully acknowledge Katja Kleemann and Tina Schneider for technical support. We thank Doron Rapaport and Chris Meisinger for antibodies; and Doron Rapaport and Till Klecker for helpful discussions.

## Data Availability

Raw and processed images have been uploaded to an OMERO repository (Burel et al., 2015) and will be released upon the final publication. In the meantime, imaging files can be requested from the corresponding author.

The mass spectrometry proteomics data have been deposited to the ProteomeXchange Consortium via the PRIDE (Deutsch et al., 2023; Perez-Riverol et al., 2025) partner repository with the dataset identifier PXD063340 and will be available upon final publication.

The code for the model in Fig 7 has been deposited on github. https://github.com/SchmollerLab/Fzo1Model

## Material and Methods

### Strain construction

All strains were haploid W303 derivates, see genotype in Supplementary Table S1. The N-terminal FLAG-AID-tag on Fzo1 was integrated using CRISPR Cas9 as described previously (Azizoğlu et al., 2023). Construction of tetracycline inducible TIR plasmid is based on (Azizoğlu et al., 2023) and estradiol inducible TIR plasmid construction is based on (Ottoz et al., 2014). Strains were otherwise constructed using standard PCR-based homologous recombination or integration of linearized plasmids. See Supplementary Table S3 for details of plasmids used for integration. All strains constructed in this study are available from the corresponding author upon request.

### Cultivation conditions

Cultures were grown to log phase in SC-Glucose (1.7 g/l yeast nitrogen base without amino acids (US Biological), 5 g/l ammonium sulfate, 50 mM potassium phthalate, pH adjusted to 5 with KOH, synthetic complete with all amino acids, see Table S2 for concentrations). TIR expression was induced by addition of anhydrotetracycline (aTC, 0.2 mg/mL Stock, dissolved in 100 % ethanol) to a final concentration of 50 ng/mL for 2 h. Cultures were diluted to OD_600_ = 0.1 and depletion was induced by addition of 5 µL 5-Ph-IAA (2-(5-Phenyl-1H-indol-3-yl)acetic acid, 100 mM Stock, dissolved in DMSO) to a final concentration of 2 µM. Cultures were diluted every 3 h and OD_600_ was determined every hour. Samples were withdrawn for DNA isolation and protein extraction (see below) before dilution, frozen in liquid nitrogen and stored at -70 °C. Experiments in which mitochondrial translation was inhibited were performed in a Δ*pdr5* background to ensure efficient translation inhibition (Roussou et al., 2024). Cultures were grown to log phase and treated with 2 mg/mL Chloramphenicol (CAP) and either untreated or Fzo1-depleted (see above).

### Microfluidic cultivation

In preparation for live cell imaging, cells were grown over night in SC-Glucose, diluted 1:50 the next morning and grown for 6 h. For the late time frame after Fzo1 depletion (8 – 18 h and 9 – 24 h), cells were grown to log phase over night, treated with 50 ng/mL aTC or 200 nM β-estradiol (dissolved at 1 mM in 100 % ethanol) for 2 h to induce TIR expression. Cells were then diluted to OD_600_ = 0.01, aTC and 5-Ph-IAA (2 µM final) were added and cells were treated for the respective time. Cells were sonicated at low power for 3 s and loaded onto a commercial microfluidics system (Y04C-02 plates, CellASIC ONIX2 system, Merck). SC-Glucose medium, with either no addition for untreated cells or supplemented with 100 ng/mL aTC or 800 nM β-estradiol (due to the hydrophobicity of the PDMS higher concentrations are required) and 2 µM 5-Ph-IAA for depletion, was supplied with a pressure of 3 psi. Cells were grown inside the microfluidic chamber for the duration of 1.5 h before imaging start. The temperature was kept constant at 30°C using an incubator chamber surrounding the imaging system (Okolab Cage Incubator, Okolab USA INC, San Bruno, CA).

### Epifluorescence microscopy

Epifluorescence microscopy was used for most imaging experiments. These were performed on a Nikon Ti2 inverted epifluorescence microscope (Nikon Instruments, Japan) with a Lumencor SPECTRA X light engine (Lumencor, Beaverton, USA), a Photometrics Prime 95 (Teledyne Photometrics, USA) backilluminated sCMOS camera. The system was programmed and controlled by the Nikon software NIS Elements. Focus was maintained using the Nikon “Perfect Focus System.” The time-lapse images were taken using a Nikon PlanApo oil-immersion 100× objective (NA = 1.45) with a frequency of 8 min for only preSu9-mCardinal experiments and every 12 min for experiments in which two fluorophores were imaged. 9 z-slices were acquired with a step size of 0.45 µm. See Supplementary Table S4 for optical filters and Supplementary Table S5 for exposure settings. For all fluorophores and tagged proteins, we checked for absence of phototoxicity (Cuny et al., 2022) by comparing growth rates or MMP at varying exposures and frame rates.

### Confocal microscopy

Confocal live-cell imaging was used for Figures 1B,C, S3A,B, S4F and carried out on a Zeiss LSM 800 microscope equipped with an Axiocam 506 camera, and using a 63×/1.4 NA oil DIC objective. The Incubator XLmulti S1, Pecon was used to maintain a temperature of 30 °C. mCardinal was imaged with an excitation wavelength of 561 nm and detected with an emission between 610 and 700 nm. Bright-field images were taken using the transmitted light detector (T-PMT). 15 z-slices with a step size of 0.5 µm were taken every 30 seconds or every 3 minutes. For control experiment in Figure S3A,B 35 z-slices with a step size of 0.23 were taken at the respective time points.

### Image analysis and data processing

Images were recorded at 12-bit gray scale and then converted to tiffs in Cell-ACDC (Padovani et al., 2022). Cell segmentation was performed in Cell-ACDC using YeaZ (Dietler et al., 2020), minimum area: 10 pixels, minimum solidity 0.5, maximal elongation 3.0). Each time frame was then visually inspected and any segmentation and tracking errors corrected in Cell-ACDC. Cell volumes were determined automatically in Cell-ACDC as described in (Padovani et al., 2022). The median background fluorescence (cell-free area) for each image was determined and subtracted from the signal at each time point. Cell cycle states and the mother-bud connections were annotated manually in Cell-ACDC which allows us to extract mother-daughter relationships and generation numbers. Mitochondria were segmented in 3D using SpotMAX (Padovani et al., 2024) based on mitochondrially localized preSu9-mCardinal. For pre-processing in SpotMAX we used a Gaussian filter with sigma 0.7 followed by a Sato filter with two sigmas 1.0 and 1.5 to enhance network-like structures (Sato et al., 1998). The pre-processed images are then segmented by SpotMAX using Otsu thresholding (Otsu, 1979). Mitochondrial amounts of imported preSu9- mCardinal and Atp6-Neongreen were calculated by Cell-ACDC as the amount of mCardinal or Neongreen within the 3-dimensional mitochondrial segmentation mask. The amounts are calculated as follows: amount = (mean_obj - median_background)*area_obj. Cytosolic protein amounts were determined as follows: A 3D mask was generated from the cell segmentation by stacking the middle slice in a cylinder. From this, the mitochondrial segmentation was subtracted to obtain an approximation of the cytosol. Mitochondrial membrane potential (MMP) using the MitoLoc reporter modified from (Vowinckel et al., 2015) was calculated as follows:

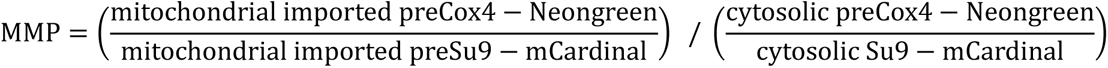

### SDS-PAGE and Western Blot

Pellets were frozen in liquid nitrogen and stored at – 70 °C. Lysates were prepared with a FastPrep® shaking three times 40 s at 6 m/s with a 1-minute break in between each cycle using lysis buffer (50 mM Tris-HCl pH 8.0, 150 mM NaCl, 5 mM EDTA, 1 % Tergitol) supplemented with 2 × EDTA-free protease inhibitor (GoldBio). 30 – 50 µg total protein were loaded on 8, 10 or 14 % SDS polyacrylamide (29:1 Bio-Rad) gels. SDS gels were blotted using a commercial wet transfer system (Bio-Rad). For detection of 3xFLAG anti-FLAG M2 antibody (Sigma, product number: F1804) (1:5000 in milk powder) and anti-mouse IgG (H+L), HRP conjugate (Promega, W4021) were used. For detection of Cox2 a custom polyclonal rabbit anti-Cox2 antibody (Singhal et al., 2017) (1:2000 in milk powder) and anti-rabbit IgG (H+L), HRP conjugate (Promega, W4011) were used. Luminescence was imaged on a Licor Odyssey FC. Bands were quantified using the image analysis software Image Studio Lite Ver 5.2.5 (Li- COR https://licor.app.box.com/s/4hrk823vov7vittqjg3onj51tb0wbo6w).

### DNA quantitative PCR (qPCR)

To isolate complete genomic DNA (gDNA), phenol-chloroform-isoamyl alcohol (PCI) extraction was performed. For this, the pellets were first dissolved in 200 μL DNA extraction buffer (pH 8.0) and transferred into a 1.5 mL safe-lock tube containing 200 μL PCI and approximately 300 µg of glass beads. The cells were mechanically disrupted by vortexing for 20 – 30s, incubated for 5 min. Then, 200 µL TE buffer were added, samples were inverted a few times and centrifuged at 13300 rpm for 5 min. 250 µL aqueous phase was then transferred to a new reaction tube containing 1 mL of 100% EtOH to precipitate the gDNA. The mixture was then centrifuged again for 5 min at 13300 rpm and the supernatant was discarded. The gDNA pellet was washed with 70% EtOH, centrifuged at 13300 rpm for 5 min, and the supernatant was again discarded. The pellet was dried at 40°C for 20 min and dissolved in 50 μL of nuclease-free water. DNA for quantitative PCR was subjected to RNase A digestion to remove RNA residues. Therefore, the solubilized pellet was treated with 1 mg/mL RNase A (ribonuclease A, 10 mg/mL, 50 U/mg, DNase-free, Roche) and incubated at 37°C for 30 min. To inactivate RNase A, DNA extraction buffer and PCI were added, and the extraction steps were repeated. One microliter of the sample was loaded on an NP80 spectrophotometer and the concentration was determined by absorbance at 260 nm. For qPCR, the samples were diluted to 0.25 ng/µL and 1 ng of DNA was used in 10 µL total reaction volume.

qPCR was performed on a QuantStudio 7 Flex Real-Time PCR system (Applied Biosystems, Thermo Fisher Scientific, Waltham, USA) in 384 multiwell plates. For amplification, a DNA-binding fluorescent dye (FastStart Universal SYBR Green Master (Rox), Roche) and specific primers for the nuclear DNA gene *ACT1* and the mtDNA genes *COX2*, *COX3*, and *COB* (Supplementary Table S6) were used. For calculation of the mtDNA fold change, the average Ct of 3 technical replicates was used. Technical replicates were excluded if the standard deviation was higher than 0.5. The average of the nuclear control gene *ACT1* was then subtracted for each gene from the average Ct for each sample. Each gene was normalized on the 0 h Control sample. The fold change was then calculated by 2^-(ΔΔCT)^ The average change of mtDNA encoded genes is presented as the average of the three mitochondrial encoded genes *COX1*, *COX2* and *COB*.

### Electron microscopy

For electron microscopy, cells were grown as described in cultivation conditions. For 21 h samples, cells were diluted to a lower OD_600_ after the 9 h timepoint and grown until 21 h to log phase. Microwave-assisted preparation and analysis of electron microscopy samples was performed as published recently (Mayer et al., 2024).

### Proteomics

Prior to mass spectrometric analysis, proteins were precipitated using acetone precipitation. Briefly, protein lysates were mixed with eight volumes of acetone and one volume of methanol. After incubation overnight at -20 °C, the protein was pelleted by centrifugation at 2 500 xg for 20 minutes at 4 °C and washed once with 80 % acetone. Protein pellets were air-dried and resuspended in denaturation buffer. Subsequently, protein quantification was performed using Bradford assay. Absorbance was measured at 595 nm and concentrations were interpolated using a standard curve of BSA. For in-solution digestion, 10 µg of each sample were diluted to a concentration of 1 mg/mL with denaturation buffer. Cysteine bonds were reduced by incubation with 1 mM DTT for 1 h at RT before carbamidomethylation by incubation with 550 nM IAA for 1 h at RT in the dark. Predigestion was performed with LysC for 3 h at RT, followed by digestion with Trypsin over night at RT. The reaction was quenched by addition of 1 % TFA and peptides were purified using StageTips (Rappsilber et al., 2007).

Mass spectrometric analysis was performed using an Orbitrap Exploris 480 online coupled to an EASY nLC-1200 system (Thermo Fisher Scientific). Separation of peptides was performed using a 20 cm HPLC column with 75 µm inner diameter (CoAnnTech) in-house packed with 1.9 µm ReproSil-Pur C18 -AQ silica beads (Dr Maisch GmbH) and elution by solvent B in a 60-minute linear gradient from 5 – 33 % at a flow rate of 200 nL/min. 30-minute wash runs were performed after each sample to minimize carry-over. Ionization was achieved by ESI and the mass spectrometer was operated in positive ion mode controlled by XCalibur (Thermo Fisher Scientific).

Spectra were acquired with a scan range of 350 – 950 m/z and a resolution on the MS1 level of 60 000. MS2 scans were performed using DIA isolation windows of 8 m/z with an overlap of 1 m/z (75 isolation windows), HCD collision energy of 30 % and resolution of 15 000.

Measurements were performed with a scan range of 350 – 950 m/z and a resolution on the MS1 level of 60 000. MS2 scans were performed using DIA isolation windows of 8 m/z with an overlap of 1 m/z (75 isolation windows), a HCD collision energy of 30 % and a resolution level of 15 000.

Raw files were processed using Spectronaut version 18 in directDIA+ mode using default settings (Bruderer et al., 2017). Spectra were predicted using a Uniprot Saccharomyces cerevisiae database with the TIR protein added by hand (53 563 entries, downloaded on 2022/12/16). The data were exported using the protein pivot report using PG.ProteinGroups as a unique protein id and the PG.Quantity as a quantification column. Downstream data analysis was performed using Perseus version 2.0.5.0 (Tyanova et al., 2016) and Microsoft Excel. Protein functions and compartments were annotated using Gene Ontology Biological Processes and Gene Ontology Cellular Compartment databases as well as Saccharomyces Genome Database for mitochondrial sub-compartments all downloaded on 2023/12/12. Statistical analysis of proteomes of 0 h vs. Control and 21 h vs. *Δfzo1* was done by Student’s t-test. Hierarchical clustering of important processes altered by Fzo1 depletion was done after filtering for the selected GO terms using z-scores while preserving the order of rows as well as columns (supervised hierarchical clustering).

### Model construction and simulation

Model simulations were performed in Matlab R2017a. Each simulation was initiated with a population of 10000 cells, each of which contained 16 nucleoids and each about to divide. At division, each cell distributed its nucleoids between the mother cell and the bud with a ratio that was randomly drawn from a log-normal distribution and rounded to integers. The parameters for the log-normal distribution were different depending on whether wild-type or Fzo1-depleted cells were simulated and were determined from experiments (Fig S7A). Specifically, for wild-type we used experimental measurements of the entire experiment, while for Fzo1-depleted cells we included data from 3-9 h after onset of depletion.

To keep the number of cells constant throughout the simulation, after cell division, the number of total cells was reduced by 50% to 10000 by random selection. To simulate mtDNA synthesis during the cell cycle, the number of nucleoids in each cell was increased by a factor that depends on whether the nucleoid number in the cell is very low (n<4), low (4=<n<7), medium (7=<n<10), high (10=<n<13), or very high (n>13). The corresponding parameters were again determined from experiments and were different for wild-type (entire experiment) and Fzo1-depleted cells (3-9 h after onset of depletion). Specifically, to categorize cells from the experiment, we measured the Atp6 amount at birth. The cells were then categorized accordingly: for example, cells with less than 3.5/8 of the mean were categorized as ‘very low’. This was done because the mean amount in the population was assumed to correspond to cells that are born with 8 nucleoids. Note that we chose the ranges of these categories for the continuous values obtained from experiments in such a way that, through nearest-integer rounding, they most appropriately reflect the discrete nucleoid values in the model. We then measured the mean relative Atp6 synthesis rate across all cells in the category (Fig S7B). After simulating synthesis, nucleoid numbers were again rounded to integers. Continuing with the simulation of the next cell division, the entire simulation was repeated 100 times.

Note that we did not consider in the simulation any variability in cell size or cell cycle duration. To test whether our simulation results are robust to changes in the initial distribution of nucleoids, we repeated it with normally distributed nucleoid numbers at time 0, mimicking the experimentally determined distribution at cell birth, and found no qualitative differences.

---

**Revision Fig.**
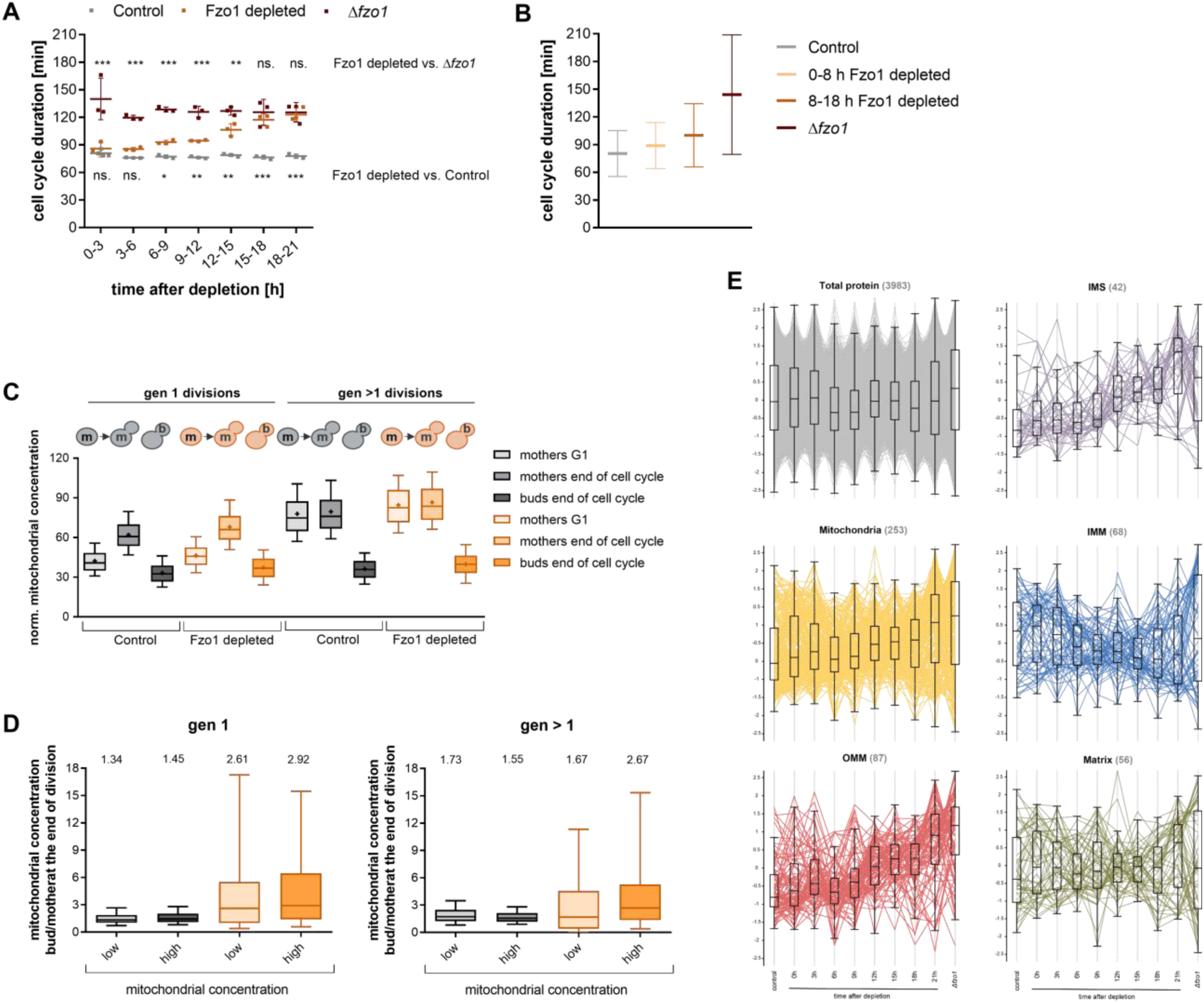
A) Unnormalized doubling times, based on same data as shown in Fig 1D. Statistical significance was determined using two-way Anova comparing the Control and Δfzo1 each to the Fzo1 depletion, assuming sphericity (equal variability between the differences). B) Cell cycle durations from live-cell microscopy. Mean with SD is shown. C) Cell volumes of first and higher generation divisions corresponding to the same cells shown in Fig 4F. D) Mitochondrial concentration ratios of bud/mothers at the end of division based on the mitochondrial concentration at the end of G1. Grouping is based on the respective Control comparison. gen 1 (Control): n (low) = 272, n (high) = 272, gen 1 (Fzo1 depleted): n (low) = 164, n (high) = 425, gen > 1 (Control): n (low) = 353, n (high) = 353, gen > 1 (Fzo1 depleted): n (low) = 473, n (high) = 262. E) z-scores of proteomic measurements of the total protein and the different mitochondrial sub-compartments.

## References

Abeliovich, H. (2023). Mitophagy in yeast: known unknowns and unknown unknowns. Biochem J, 480(20), 1639–1657. 10.1042/bcj20230279

Adams, W. C., Chen, Y.-H., Kratchmarov, R., Yen, B., Nish, S. A., Lin, W.-H. W., Rothman, N. J., Luchsinger, L. L., Klein, U., Busslinger, M., Rathmell, J. C., Snoeck, H.-W., & Reiner, S. L. (2016). Anabolism-Associated Mitochondrial Stasis Driving Lymphocyte Differentiation over Self-Renewal. Cell Reports, 17(12), 3142–3152. 10.1016/j.celrep.2016.11.065

Alexander, C., Votruba, M., Pesch, U. E. A., Thiselton, D. L., Mayer, S., Moore, A., Rodriguez, M., Kellner, U., Leo-Kottler, B., Auburger, G., Bhattacharya, S. S., & Wissinger, B. (2000). OPA1, encoding a dynamin-related GTPase, is mutated in autosomal dominant optic atrophy linked to chromosome 3q28. Nature Genetics, 26(2), 211–215. 10.1038/79944

Azizoğlu, A., Loureiro, C., Venetz, J., & Brent, R. (2023). Autorepression-Based Conditional Gene Expression System in Yeast for Variation-Suppressed Control of Protein Dosage. Curr Protoc, 3(1), e647. 10.1002/cpz1.647

Böckler, S., Chelius, X., Hock, N., Klecker, T., Wolter, M., Weiss, M., Braun, R. J., & Westermann, B. (2017). Fusion, fission, and transport control asymmetric inheritance of mitochondria and protein aggregates. The Journal of cell biology, 216(8), 2481–2498. 10.1083/jcb.201611197

Bruderer, R., Bernhardt, O. M., Gandhi, T., Xuan, Y., Sondermann, J., Schmidt, M., Gomez-Varela, D., & Reiter, L. (2017). Optimization of Experimental Parameters in Data-Independent Mass Spectrometry Significantly Increases Depth and Reproducibility of Results. Mol Cell Proteomics, 16(12), 2296–2309. 10.1074/mcp.RA117.000314

Burel, J. M., Besson, S., Blackburn, C., Carroll, M., Ferguson, R. K., Flynn, H., Gillen, K., Leigh, R., Li, S., Lindner, D., Linkert, M., Moore, W. J., Ramalingam, B., Rozbicki, E., Tarkowska, A., Walczysko, P., Allan, C., Moore, J., & Swedlow, J. R. (2015). Publishing and sharing multi-dimensional image data with OMERO. Mamm Genome, 26(9-10), 441–447. 10.1007/s00335-015-9587-6

Chacko, L. A., Nakaoka, H., Morris, R., Marshall, W., & Ananthanarayanan, V. (2025). Mitochondrial function regulates cell growth kinetics to actively maintain mitochondrial homeostasis. bioRxiv, 2025.2003.2031.646474. 10.1101/2025.03.31.646474

Contamine, V., & Picard, M. (2000). Maintenance and integrity of the mitochondrial genome: a plethora of nuclear genes in the budding yeast. Microbiol Mol Biol Rev, 64(2), 281–315. 10.1128/mmbr.64.2.281-315.2000

Cuny, A. P., Schlottmann, F. P., Ewald, J. C., Pelet, S., & Schmoller, K. M. (2022). Live cell microscopy: From image to insight. Biophys Rev (Melville), 3(2), 021302. 10.1063/5.0082799

Dagsgaard, C., Taylor, L. E., O’Brien, K. M., & Poyton, R. O. (2001). Effects of anoxia and the mitochondrion on expression of aerobic nuclear COX genes in yeast: evidence for a signaling pathway from the mitochondrial genome to the nucleus. J Biol Chem, 276(10), 7593–7601. 10.1074/jbc.M009180200

Delettre, C., Lenaers, G., Griffoin, J.-M., Gigarel, N., Lorenzo, C., Belenguer, P., Pelloquin, L., Grosgeorge, J., Turc-Carel, C., Perret, E., Astarie-Dequeker, C., Lasquellec, L., Arnaud, B., Ducommun, B., Kaplan, J., & Hamel, C. P. (2000). Nuclear gene OPA1, encoding a mitochondrial dynamin-related protein, is mutated in dominant optic atrophy. Nature Genetics, 26(2), 207–210. 10.1038/79936

Deutsch, E. W., Bandeira, N., Perez-Riverol, Y., Sharma, V., Carver, J. J., Mendoza, L., Kundu, D. J., Wang, S., Bandla, C., Kamatchinathan, S., Hewapathirana, S., Pullman, B. S., Wertz, J., Sun, Z., Kawano, S., Okuda, S., Watanabe, Y., MacLean, B., MacCoss, M. J., … Vizcaíno, J. A. (2023). The ProteomeXchange consortium at 10 years: 2023 update. Nucleic Acids Res, 51(D1), D1539–d1548. 10.1093/nar/gkac1040

Di Bartolomeo, F., Malina, C., Campbell, K., Mormino, M., Fuchs, J., Vorontsov, E., Gustafsson, C. M., & Nielsen, J. (2020). Absolute yeast mitochondrial proteome quantification reveals trade-off between biosynthesis and energy generation during diauxic shift. Proc Natl Acad Sci U S A, 117(13), 7524–7535. 10.1073/pnas.1918216117

Dietler, N., Minder, M., Gligorovski, V., Economou, A. M., Joly, D. A. H. L., Sadeghi, A., Chan, C. H. M., Koziński, M., Weigert, M., Bitbol, A.-F., & Rahi, S. J. (2020). A convolutional neural network segments yeast microscopy images with high accuracy. Nature Communications, 11(1), 5723. 10.1038/s41467-020-19557-4

Friedman, J. R., & Nunnari, J. (2014). Mitochondrial form and function. Nature, 505(7483), 335–343. 10.1038/nature12985

Gorospe, C. M., Carvalho, G., Herrera Curbelo, A., Marchhart, L., Mendes, I. C., Niedźwiecka, K., & Wanrooij, P. H. (2023). Mitochondrial membrane potential acts as a retrograde signal to regulate cell cycle progression. Life Science Alliance, 6(12), e202302091. 10.26508/lsa.202302091

Hermann, G. J., Thatcher, J. W., Mills, J. P., Hales, K. G., Fuller, M. T., Nunnari, J., & Shaw, J. M. (1998). Mitochondrial fusion in yeast requires the transmembrane GTPase Fzo1p. The Journal of cell biology, 143(2), 359–373. 10.1083/jcb.143.2.359

Hinge, A., He, J., Bartram, J., Javier, J., Xu, J., Fjellman, E., Sesaki, H., Li, T., Yu, J., Wunderlich, M., Mulloy, J., Kofron, M., Salomonis, N., Grimes, H. L., & Filippi, M.-D. (2020). Asymmetrically Segregated Mitochondria Provide Cellular Memory of Hematopoietic Stem Cell Replicative History and Drive HSC Attrition. Cell Stem Cell, 26(3), 420–430.e426. 10.1016/j.stem.2020.01.016

Jajoo, R., Jung, Y., Huh, D., Viana, M. P., Rafelski, S. M., Springer, M., & Paulsson, J. (2016). Accurate concentration control of mitochondria and nucleoids. Science, 351(6269), 169–172. 10.1126/science.aaa8714

Jakobs, S., Martini, N., Schauss, A. C., Egner, A., Westermann, B., & Hell, S. W. (2003). Spatial and temporal dynamics of budding yeast mitochondria lacking the division component Fis1p. Journal of Cell Science, 116(10), 2005–2014. 10.1242/jcs.00423

Katajisto, P., Döhla, J., Chaffer, C. L., Pentinmikko, N., Marjanovic, N., Iqbal, S., Zoncu, R., Chen, W., Weinberg, R. A., & Sabatini, D. M. (2015). Stem cells. Asymmetric apportioning of aged mitochondria between daughter cells is required for stemness. Science, 348(6232), 340–343. 10.1126/science.1260384

Khan, A. H., Gu, X., Patel, R. J., Chuphal, P., Viana, M. P., Brown, A. I., Zid, B. M., & Tsuboi, T. (2024). Mitochondrial protein heterogeneity stems from the stochastic nature of co-translational protein targeting in cell senescence. Nature Communications, 15(1), 8274. 10.1038/s41467-024-52183-y

Klecker, T., & Westermann, B. (2021). Pathways shaping the mitochondrial inner membrane. Open Biol, 11(12), 210238. 10.1098/rsob.210238

Knorre, D. A., Sokolov, S. S., Zyrina, A. N., & Severin, F. F. (2016). How do yeast sense mitochondrial dysfunction? Microb Cell, 3(11), 532–539. 10.15698/mic2016.11.537

Leonard, P. J., Rathod, P. K., & Golin, J. (1994). Loss of function mutation in the yeast multiple drug resistance gene PDR5 causes a reduction in chloramphenicol efflux. Antimicrob Agents Chemother, 38(10), 2492–2494. 10.1128/aac.38.10.2492

Liu, S., Liu, S., He, B., Li, L., Li, L., Wang, J., Cai, T., Chen, S., & Jiang, H. (2021). OXPHOS deficiency activates global adaptation pathways to maintain mitochondrial membrane potential. EMBO Rep, 22(4), e51606. 10.15252/embr.202051606

Mayer, M., Schug, C., Geimer, S., Klecker, T., & Westermann, B. (2024). Microwave-assisted preparation of yeast cells for ultrastructural analysis by electron microscopy. Microb Cell, 11, 378–386. 10.15698/mic2024.11.840

Nunnari, J., Marshall, W. F., Straight, A., Murray, A., Sedat, J. W., & Walter, P. (1997). Mitochondrial transmission during mating in Saccharomyces cerevisiae is determined by mitochondrial fusion and fission and the intramitochondrial segregation of mitochondrial DNA. Mol Biol Cell, 8(7), 1233–1242. 10.1091/mbc.8.7.1233

Osman, C., Noriega, T. R., Okreglak, V., Fung, J. C., & Walter, P. (2015). Integrity of the yeast mitochondrial genome, but not its distribution and inheritance, relies on mitochondrial fission and fusion. Proc Natl Acad Sci U S A, 112(9), E947–956. 10.1073/pnas.1501737112

Otsu, N. (1979). A Threshold Selection Method from Gray-Level Histograms. IEEE Trans. Syst. Man Cybern., 9, 62–66.

Ottoz, D. S., Rudolf, F., & Stelling, J. (2014). Inducible, tightly regulated and growth condition-independent transcription factor in Saccharomyces cerevisiae. Nucleic Acids Res, 42(17), e130. 10.1093/nar/gku616

Padovani, F., Čavka, I., Neves, A. R. R., López, C. P., Al-Refaie, N., Bolcato, L., Chatzitheodoridou, D., Chadha, Y., Su, X. A., Lengefeld, J., Cabianca, D. S., Köhler, S., & Schmoller, K. M. (2024). SpotMAX: a generalist framework for multi-dimensional automatic spot detection and quantification. bioRxiv, 2024.2010.2022.619610. 10.1101/2024.10.22.619610

Padovani, F., Mairhörmann, B., Falter-Braun, P., Lengefeld, J., & Schmoller, K. M. (2022). Segmentation, tracking and cell cycle analysis of live-cell imaging data with Cell-ACDC. BMC Biol, 20(1), 174. 10.1186/s12915-022-01372-6

Paumard, P., Vaillier, J., Coulary, B., Schaeffer, J., Soubannier, V., Mueller, D. M., Brèthes, D., di Rago, J. P., & Velours, J. (2002). The ATP synthase is involved in generating mitochondrial cristae morphology. The EMBO Journal, 21(3), 221–230-230. 10.1093/emboj/21.3.221

Perez-Riverol, Y., Bandla, C., Kundu, D. J., Kamatchinathan, S., Bai, J., Hewapathirana, S., John, N. S., Prakash, A., Walzer, M., Wang, S., & Vizcaíno, J. A. (2025). The PRIDE database at 20 years: 2025 update. Nucleic Acids Res, 53(D1), D543–d553. 10.1093/nar/gkae1011

Quintana-Cabrera, R., & Scorrano, L. (2023). Determinants and outcomes of mitochondrial dynamics. Molecular Cell, 83(6), 857–876. 10.1016/j.molcel.2023.02.012

Rafelski, S. M., Viana, M. P., Zhang, Y., Chan, Y. H., Thorn, K. S., Yam, P., Fung, J. C., Li, H., Costa Lda, F., & Marshall, W. F. (2012). Mitochondrial network size scaling in budding yeast. Science, 338(6108), 822–824. 10.1126/science.1225720

Rapaport, D., Brunner, M., Neupert, W., & Westermann, B. (1998). Fzo1p is a mitochondrial outer membrane protein essential for the biogenesis of functional mitochondria in Saccharomyces cerevisiae. J Biol Chem, 273(32), 20150–20155. 10.1074/jbc.273.32.20150

Rappsilber, J., Mann, M., & Ishihama, Y. (2007). Protocol for micro-purification, enrichment, pre-fractionation and storage of peptides for proteomics using StageTips. Nat Protoc, 2(8), 1896–1906. 10.1038/nprot.2007.261

Ray, M. W., Chen, W., Duan, C., Bravo, G., Krueger, K., Rosario, E. M., Jacob, A. A., & Lackner, L. L. (2025). The Volume of Mitochondria Inherited Impacts mtDNA Homeostasis in Budding Yeast. bioRxiv, 2025.2003.2025.645216. 10.1101/2025.03.25.645216

Roussou, R., Metzler, D., Padovani, F., Thoma, F., Schwarz, R., Shraiman, B., Schmoller, K. M., & Osman, C. (2024). Real-time assessment of mitochondrial DNA heteroplasmy dynamics at the single-cell level. The EMBO Journal, 43(22), 5340–5359-5359. 10.1038/s44318-024-00183-5

Sato, Y., Nakajima, S., Shiraga, N., Atsumi, H., Yoshida, S., Koller, T., Gerig, G., & Kikinis, R. (1998). Three-dimensional multi-scale line filter for segmentation and visualization of curvilinear structures in medical images. Medical Image Analysis, 2(2), 143–168. 10.1016/S1361-8415(98)80009-1

Seel, A., Padovani, F., Mayer, M., Finster, A., Bureik, D., Thoma, F., Osman, C., Klecker, T., & Schmoller, K. M. (2023). Regulation with cell size ensures mitochondrial DNA homeostasis during cell growth. Nat Struct Mol Biol, 30(10), 1549–1560. 10.1038/s41594-023-01091-8

Shirozu, R., Yashiroda, H., & Murata, S. (2016). Proteasome Impairment Induces Recovery of Mitochondrial Membrane Potential and an Alternative Pathway of Mitochondrial Fusion. Mol Cell Biol, 36(2), 347–362. 10.1128/mcb.00920-15

Silva Ramos, E., Motori, E., Brüser, C., Kühl, I., Yeroslaviz, A., Ruzzenente, B., Kauppila, J. H. K., Busch, J. D., Hultenby, K., Habermann, B. H., Jakobs, S., Larsson, N. G., & Mourier, A. (2019). Mitochondrial fusion is required for regulation of mitochondrial DNA replication. PLoS Genet, 15(6), e1008085. 10.1371/journal.pgen.1008085

Singhal, R. K., Kruse, C., Heidler, J., Strecker, V., Zwicker, K., Düsterwald, L., Westermann, B., Herrmann, J. M., Wittig, I., & Rapaport, D. (2017). Coi1 is a novel assembly factor of the yeast complex III-complex IV supercomplex. Mol Biol Cell, 28(20), 2609–2622. 10.1091/mbc.E17-02-0093

Stenberg, S., Li, J., Gjuvsland, A. B., Persson, K., Demitz-Helin, E., González Peña, C., Yue, J.-X., Gilchrist, C., Ärengård, T., Ghiaci, P., Larsson-Berglund, L., Zackrisson, M., Smits, S., Hallin, J., Höög, J. L., Molin, M., Liti, G., Omholt, S. W., & Warringer, J. (2022). Genetically controlled mtDNA deletions prevent ROS damage by arresting oxidative phosphorylation. eLife, 11, e76095. 10.7554/eLife.76095

Tyanova, S., Temu, T., Sinitcyn, P., Carlson, A., Hein, M. Y., Geiger, T., Mann, M., & Cox, J. (2016). The Perseus computational platform for comprehensive analysis of (prote)omics data. Nat Methods, 13(9), 731–740. 10.1038/nmeth.3901

Vőfély, R. V., Gallagher, J., Pisano, G. D., Bartlett, M., & Braybrook, S. A. (2019). Of puzzles and pavements: a quantitative exploration of leaf epidermal cell shape. New Phytologist, 221(1), 540–552. 10.1111/nph.15461

Vowinckel, J., Hartl, J., Butler, R., & Ralser, M. (2015). MitoLoc: A method for the simultaneous quantification of mitochondrial network morphology and membrane potential in single cells. Mitochondrion, 24, 77–86. 10.1016/j.mito.2015.07.001

Vowinckel, J., Hartl, J., Marx, H., Kerick, M., Runggatscher, K., Keller, M. A., Mülleder, M., Day, J., Weber, M., Rinnerthaler, M., Yu, J. S. L., Aulakh, S. K., Lehmann, A., Mattanovich, D., Timmermann, B., Zhang, N., Dunn, C. D., MacRae, J. I., Breitenbach, M., & Ralser, M. (2021). The metabolic growth limitations of petite cells lacking the mitochondrial genome. Nat Metab, 3(11), 1521–1535. 10.1038/s42255-021-00477-6

Westermann, B. (2010). Mitochondrial fusion and fission in cell life and death. Nature Reviews Molecular Cell Biology, 11(12), 872–884. 10.1038/nrm3013

Wisniewski, B. T., Casler, J. C., & Lackner, L. L. (2024). Significantly reduced, but balanced, rates of mitochondrial fission and fusion are sufficient to maintain the integrity of yeast mitochondrial DNA. Molecular Biology of the Cell, 35(12), br25. 10.1091/mbc.E24-07-0306

Yesbolatova, A., Saito, Y., Kitamoto, N., Makino-Itou, H., Ajima, R., Nakano, R., Nakaoka, H., Fukui, K., Gamo, K., Tominari, Y., Takeuchi, H., Saga, Y., Hayashi, K. I., & Kanemaki, M. T. (2020). The auxin-inducible degron 2 technology provides sharp degradation control in yeast, mammalian cells, and mice. Nat Commun, 11(1), 5701. 10.1038/s41467-020-19532-z

Züchner, S., Mersiyanova, I. V., Muglia, M., Bissar-Tadmouri, N., Rochelle, J., Dadali, E. L., Zappia, M., Nelis, E., Patitucci, A., Senderek, J., Parman, Y., Evgrafov, O., Jonghe, P. D., Takahashi, Y., Tsuji, S., Pericak-Vance, M. A., Quattrone, A., Battologlu, E., Polyakov, A. V., … Vance, J. M. (2004). Mutations in the mitochondrial GTPase mitofusin 2 cause Charcot-Marie-Tooth neuropathy type 2A. Nature Genetics, 36(5), 449–451. 10.1038/ng1341

