## Supplementary Material for "When mitochondria fall apart: Unbalanced mitochondrial segregation triggers loss of mtDNA in the absence of mitochondrial fusion"

**Supplementary Tables:**

Table S1: Strains used in this study.

Table S2: Components of synthetic complete (SC)

Table S3: Plasmids used in this study.

Table S4: Optical filters for microscopy.

Table S5: Exposure settings and intensities.

Table S6: Primers used for DNA-qPCR

Table S7: Proteomics data (separate file)

**Supplementary Figures**

Figure S1: Control experiments on growth and mitochondrial morphology.

Figure S2: Respiratory chain proteins and mitochondrial ribosomal subunits are decreasing together with mitochondrial ultrastructure.

Figure S3: Additional experiments on mitochondrial membrane potential and growth

Figure S4: Additional experiments on mitochondrial distribution, quality and morphology.

Figure S5: Additional experiments on Atp6-Neongreen concentrations and Fzo1 depletion in the mitophagy mutant Δ*atg32*.

Figure S6: Additional experiments on the Atp6 fold change over the cell cycle.

Figure S7: Distribution of Atp6 between daughters and mothers and average Atp6 fold change of the individual categories used for the model.

**Supplementary References**

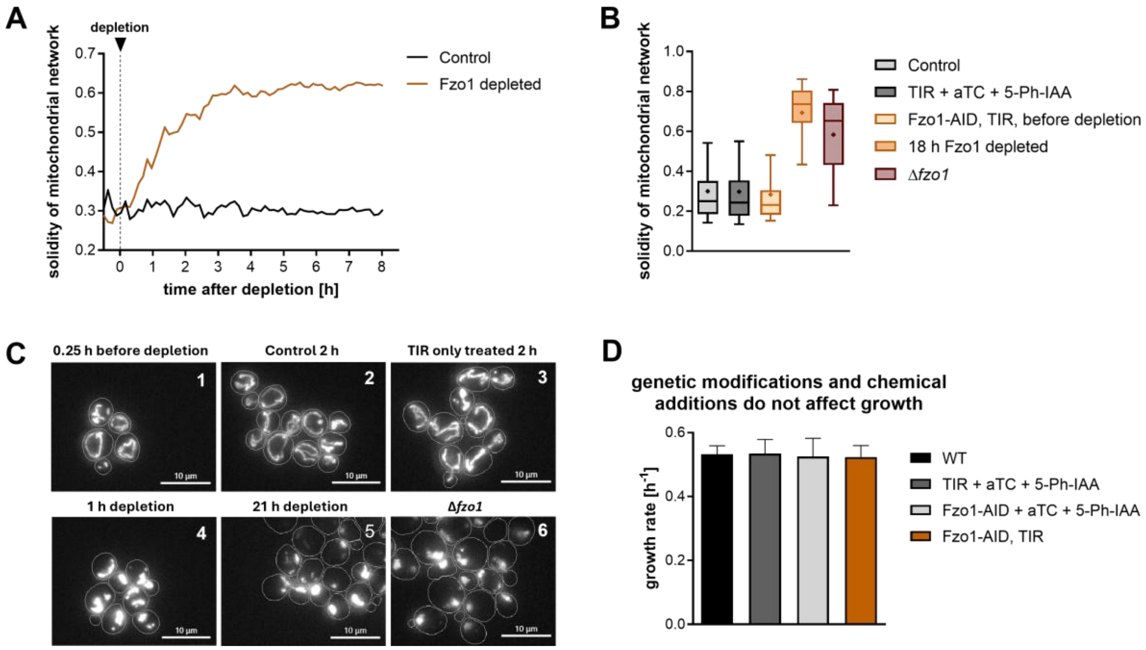


**Supplement Figure 1.** Cells were grown in synthetic complete (SC) medium containing 1 % glucose and Fzo1 depletion was induced at the indicated times. A) Solidity of the mitochondrial network from epifluorescence microscopy. Mean from three biological replicates with a total of 1694 Control and 1694 Fzo1 depleted cells. B) Solidity of Control strains from three biological replicates. Whiskers indicate 10$^{th}$ and 90$^{th}$ percentiles. Mitochondrial morphology of: Control (all 8.5 h of imaging) 26850 datapoints of 1694 cells, TIR + aTC + 5-PH-IAA (after addition of the chemicals) 17170 datapoints of 1078 cells, Fzo1-AID, TIR before depletion (0.5 h before addition of 5-Ph-IAA, aTC is added already) 207 datapoints of 57 cells, 18 h induced 1434 datapoints of 1434 cells, Δ*fzo1* (all of 10 h imaging) 43414 datapoints of 1419 cells. C) Example pictures of maximum z-projections of mitochondria visualized by pre-Su9-mCardinal. Scale bar = 10 µm. 1: Fzo1-AID, TIR + aTC, 2: Control = Fzo1-AID, TIR, 3: TIR + aTC + 5-Ph-IAA, 4: Fzo1-AID, TIR + aTC + 5-Ph-IAA, 5: Fzo1-AID, TIR + aTC + 5-Ph-IAA, 6: Δ*fzo1*. D) Growth rate of WT cells, cells with TIR treated with anhydrotetracycline (aTC) and 5-Ph-IAA, cells with FLAG-AID-Fzo1 treated with aTC and 5-Ph-IAA, and cells with TIR and FLAG-AID-Fzo1 without treatment. Mean with SD from three biological replicates is shown.

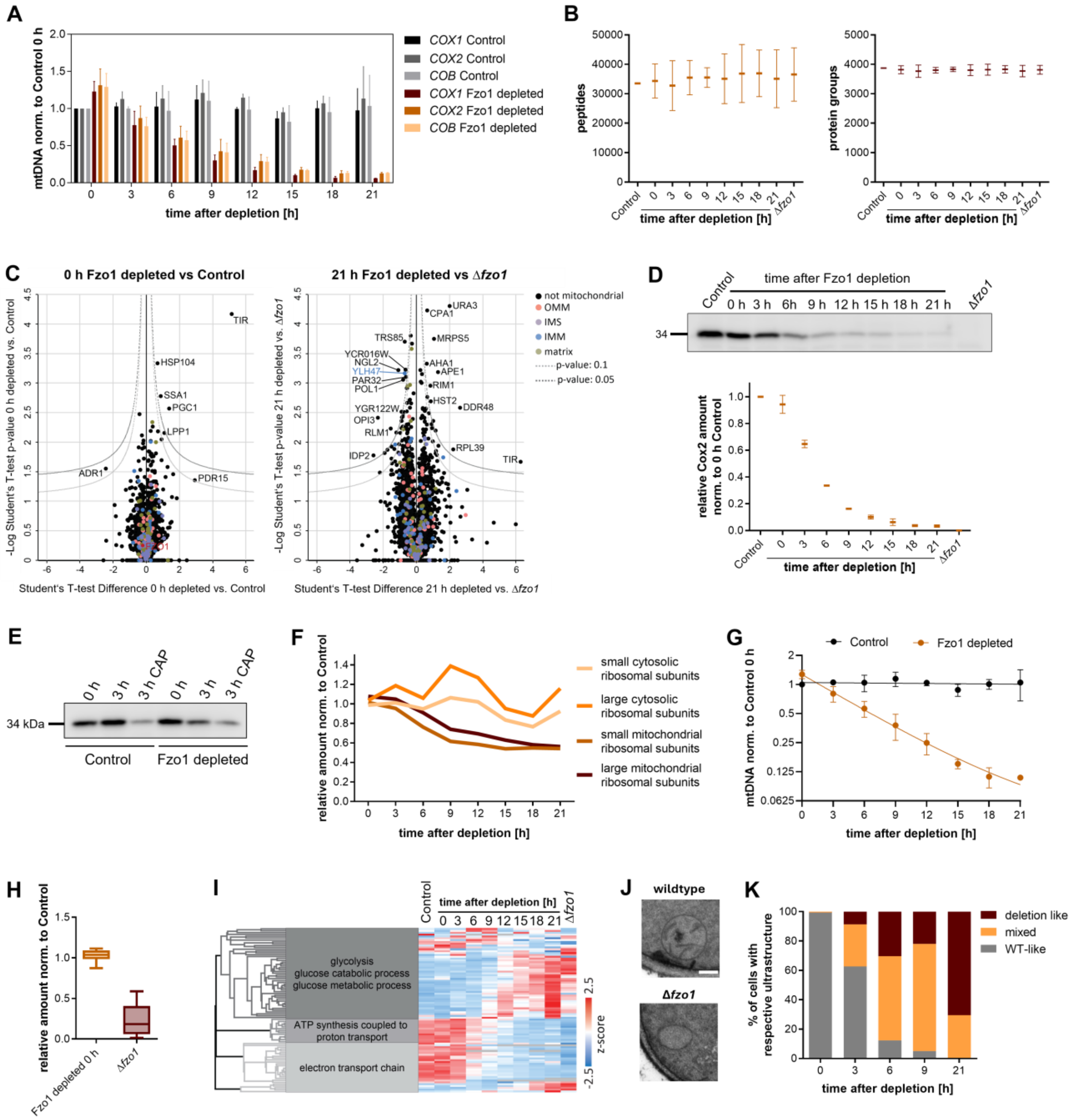


**Supplement Figure 2.** A) DNA-qPCR of individual mitochondrial genes. The level of each gene was normalized to the nuclear gene *ACT1*. Error bars indicate SD from three biological replicates. B) Numbers of peptides and protein groups detected in the mass spectrometry analysis. C) Volcano plots of proteomics measurement of 0 h depleted vs Control cells (left) and 21 h depleted vs Δ*fzo1* cells. D) Quantification of Cox2 amount by Western Blot analysis. Error bars indicate SD from three biological replicates. Example Western Blot is shown above. E) Example Western Blot of Chloramphenicol (CAP) treatment quantified in Fig 2C. F) Proteomics measurement of mitochondrial ribosomal subunits and cytosolic ribosomal subunits. The average of all measured protein groups of the respective subunits is shown G) mtDNA levels shown in Fig 2A on log2 scale. Solid line shows the one phase exponential decay with a half-life of 4.7 h for Fzo1 depleted cells. H) Proteomics measurement of nuclear encoded respiratory chain proteins of Control and Δ*fzo1* cells related to Fig 2D. I) Heat map of important processes which are altered through Fzo1 depletion. J) Example pictures of electron microscopy related to Figure 3G), scale bar = 200 nm. K) Categorization of cells based on their ultrastructure of electron microscopy of three biological replicates, related to figure 2H.

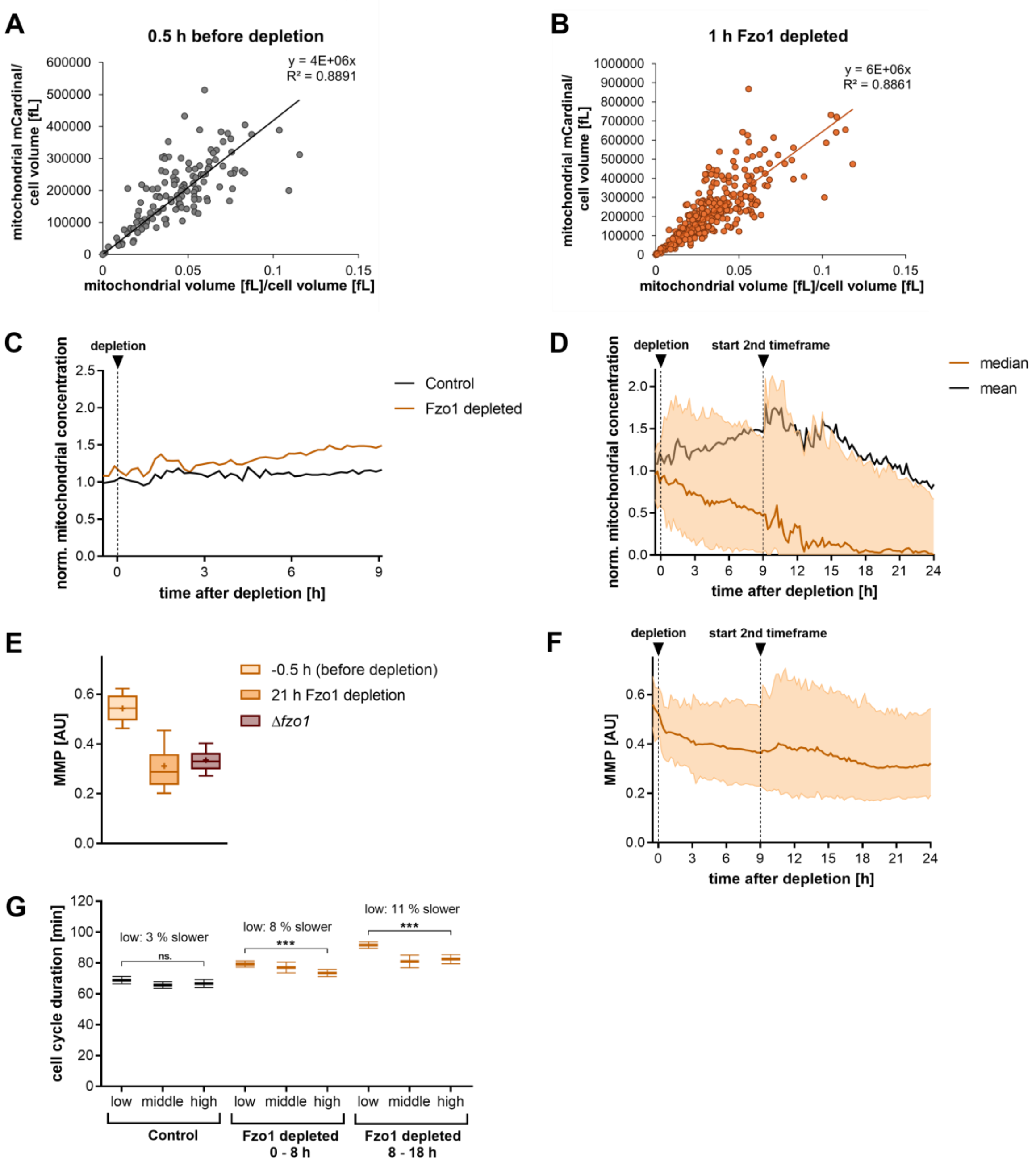


**Supplement Figure 3.** A) mitochondrial volume [fL]/cell volume [fL] vs mitochondrial mCardinal/cell volume [fL] before depletion. N = 129. determined by a 3D-reconstruction of 15 slices taken by confocal microscopy. B) mitochondrial volume [fL]/cell volume [fL] vs mitochondrial mCardinal/cell volume [fL] 1 h after Fzo1 depletion. N = 286. A) and B) total mitochondrial mCardinal signal correlates with mitochondrial volume from a 3D volume reconstruction. However, due to the morphology change after Fzo1 depletion the slope of the relationship changes, likely due to a systematic underestimation of the mitochondrial volume in fragmented aggregated mitochondria. Therefore, the total mitochondrial mCardinal is the best proxy for the total content of mitochondrial network in the cell. In the following, we thus use the term “mitochondrial concentration” for total mitochondrial mCardinal intensity per cell volume. C) Mean of the normalized mitochondrial concentrations after Fzo1 depletion. Same data as in Fig 3A. D) Normalized mitochondrial concentrations after Fzo1 depletion. Shaded areas show 25$^{th}$ and 75$^{th}$ percentiles. Cells were imaged in two time intervals: -0.5 – 9 h and 9 – 24 h after Fzo1 depletion. n (24 h Fzo1 depletion) = 2019. E) MMP of controls as described in Fig 3. -0.5 h before depletion: 184 datapoints of 67 cells, 21 h Fzo1 depletion: 718 datapoints of 718 cells, Δ*fzo1:* 25197 datapoints of 1376 cells over 10 h of imaging. F) MMP of Fzo1 depleted cells show in D). Shaded areas show 5$^{th}$ and 95$^{th}$ percentiles. n (24 h Fzo1 depletion) = 2019. G) Cell cycle duration of generation >1 with different mitochondrial concentrations. Categorization was performed based on Control cells gen >1. Images were taken every 8 min to determine cell cycle durations. Mean with 95 % Confidence interval is shown. Statistical significance was determined using a paired two-tailed t-test. *** indicate $p < 0.001$, n.s. = not significant. Control: n (low) = 233, n (middle) = 233, n (high) = 240, Fzo1 depleted 0 – 8 h: n (low) = 460, n (middle) = 98, n (high) = 232, Fzo1 depleted 8 – 18 h: n (low) = 714, n (middle) = 82, n (high) = 232.

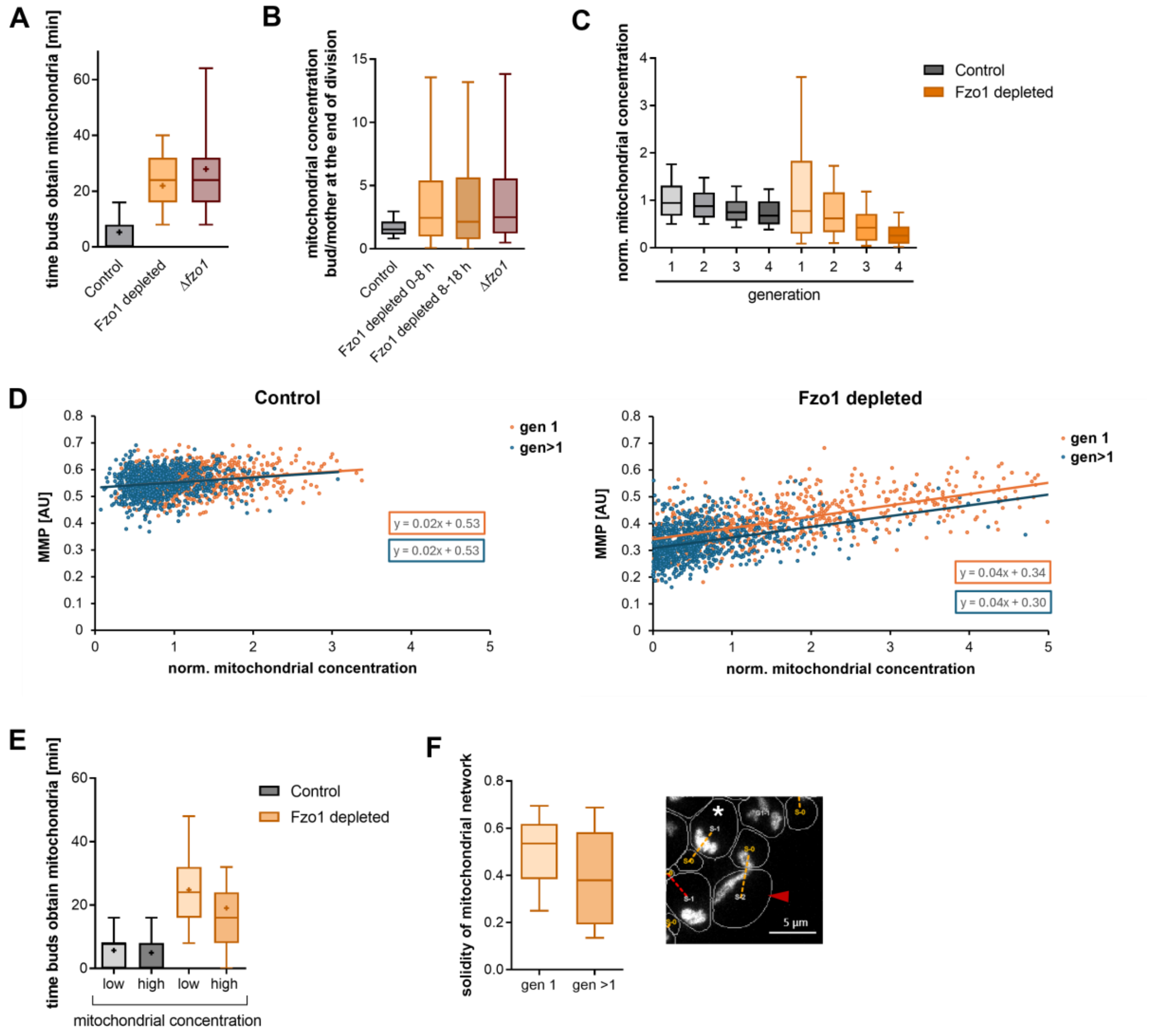


**Supplement Figure 4.** A) Time between start of budding and mitochondria are detected in the bud. Control = 1249, Fzo1 depleted = 1275, Δ*fzo1* = 1237. B) Ratio of the mitochondrial concentration (total mitochondrial mCardinal intensity per cell volume) of buds to mothers at the end of the cell cycle. n (Control) = 1250, n (Fzo1 depleted 0 – 8 h) = 1324, n (Fzo1 depleted 8 – 18 h) = 1410, n (Δ*fzo1*) = 1226. C) Mitochondrial concentrations of cells shown in Fig 4I. D) Mitochondrial concentration against MMP of mothers of gen 1 or gen > 1 for Control and Fzo1 depleted cells (0 – 9 h following Fzo1 depletion). Control: n (gen 1) = 687, n (gen >1) = 943. Fzo1 depleted: n (gen 1) = 745, n (gen >1) = 1024. E) The start of budding and when mitochondria are first detected in the bud of mothers with a low or high mitochondrial concentration. Time is from start of budding until mitochondria are detected in the bud. The median mitochondrial concentration of Control cells at G1 was used to determine low and high categories. Control: n (low) = 625, n (high) = 624, Fzo1 depleted: n (low) = 638, n (high) = 637. F) Solidity of gen 1 and gen > 1 mothers throughout their cell cycle is shown. Mitochondrial morphology was determined for every time in the cell cycle. n > 11000 datapoints of at least 257 cells each. Red arrow on the example image shows a gen > 1 mother, white star shows a first generation mother. Scale bar = 5 µm. The differences in mitochondrial morphology is likely one factor which contributes to the lower inheritance of mitochondria to buds in higher generations. In higher generation cell cycles the mitochondrial network of the mothers more likely exhibits strings as observed by a lower solidity than first generation divisions. This indicates that in higher generation mothers mitochondria are likely more strongly tethered to the cell cortex than in first generation divisions, which possibly contributes to a lower inheritance of mitochondria to buds. Of note, these are only short morphological changes and are not a sign of incomplete Fzo1 depletion in these mitochondria.

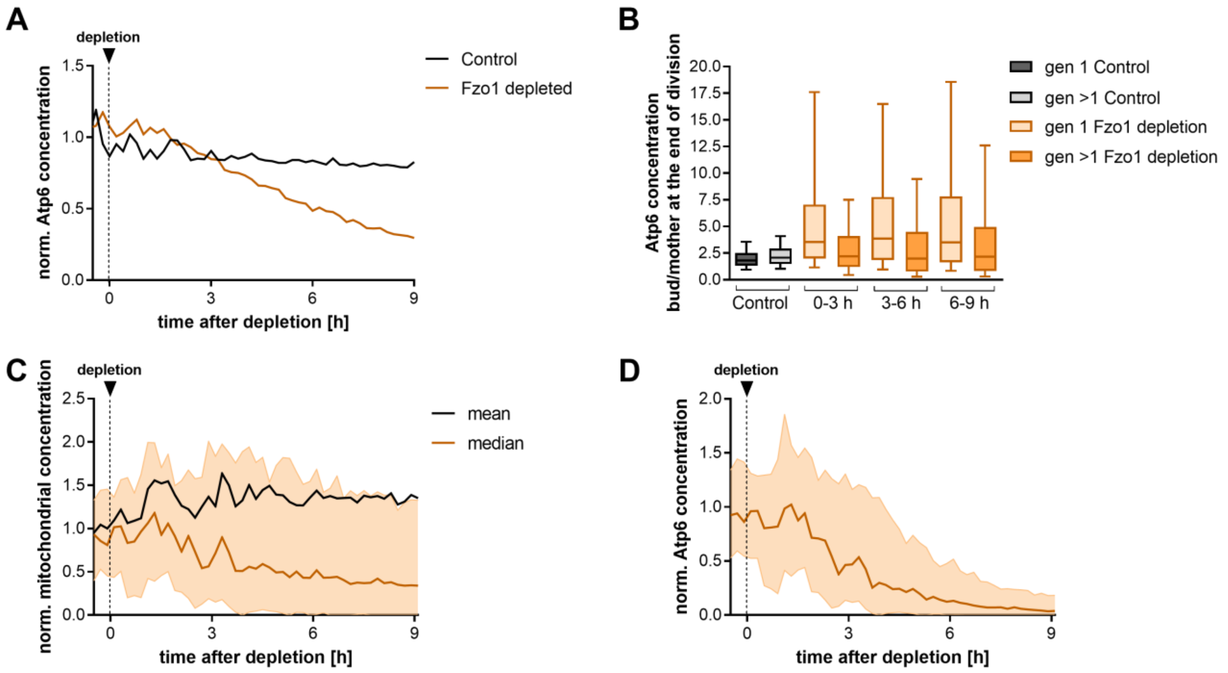


**Supplement Figure 5.** A) Atp6 concentration after Fzo1 depletion. Same data as in main Fig 5A. B) Ratio of the Atp6 concentration between buds and mothers at the end of division of cells shown in Fig 5C). C) Mitochondrial concentration after Fzo1 depletion in Δ*atg32* cells which are unable to perform mitophagy. Median with 25$^{th}$ and 75$^{th}$ percentiles is shown in orange, mean is shown in black. Analysis of 1909 cells at 9 h from two biological replicates is shown. See Fig 3A and S3A for comparison of Fzo1 depletion in a wildtype background. D) Atp6-Neongreen concentration of Δ*atg32* cells after Fzo1 depletion. Median with 25$^{th}$ and 75$^{th}$ percentiles of 1909 cells at 9 h from two biological replicates is shown. See Fig 5A for comparison of Fzo1 depletion in a wildtype background.

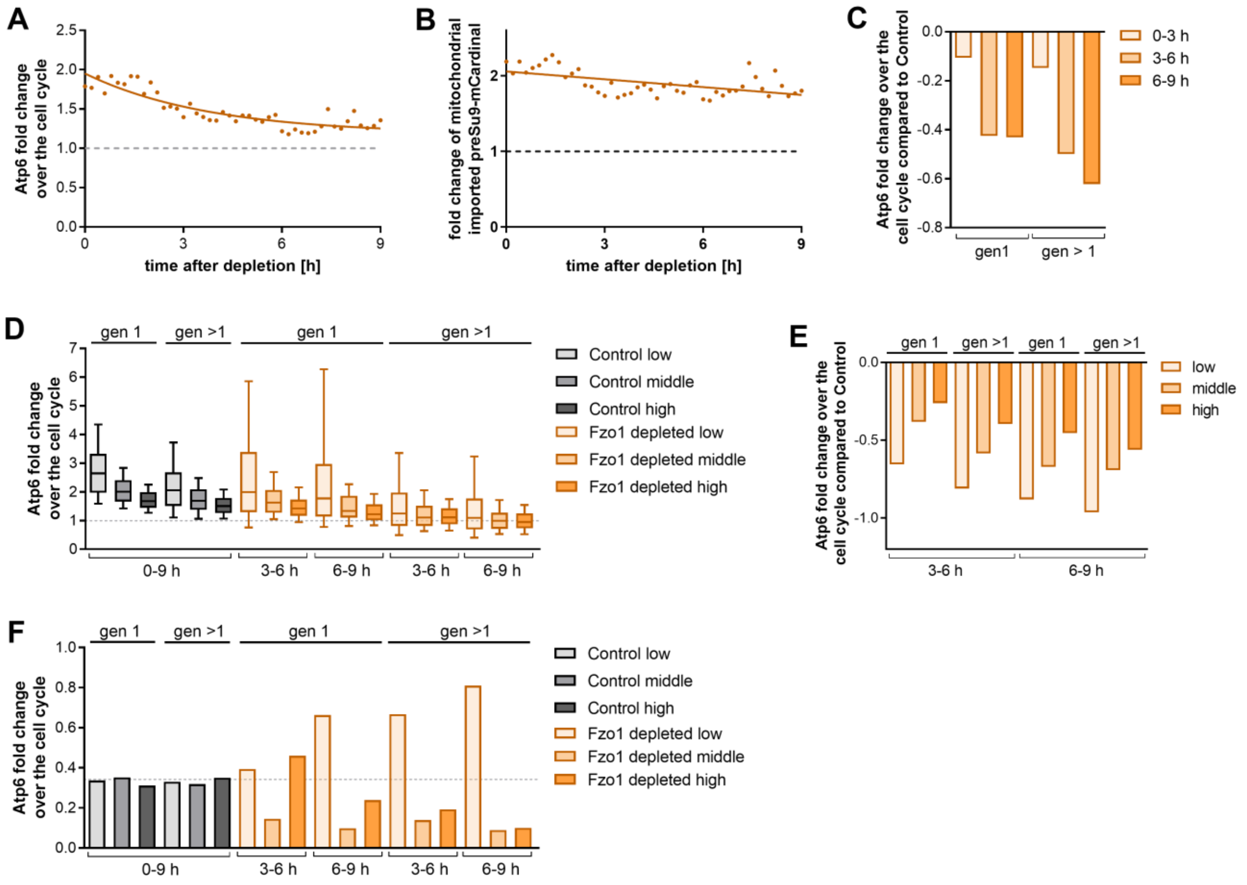


**Supplement Figure 6.** A) Fold change of Atp6 over the cell cycle of all generations after Fzo1 depletion related to Fig 6B. 9013 cell divisions were analyzed. B) Fold change of mitochondrial imported preSu9-mCardinal over the cell cycle of all generations after Fzo1 depletion as in A). C) Difference of the median fold change of Atp6 over the cell cycle compared to Control cells of gen 1 and gen >1 divisions, related to Fig 6C. D) Fold change of Atp6 over the cell cycle of first and higher generation divisions of mothers with low, middle, or high Atp6 content at the beginning of the cell cycle. At least 79 cell cycles were analyzed for each group. E) Difference of the median Atp6 fold change of Fzo1 depleted cells compared to the respective Control group of cells shown in D). F) Fraction of the population of cells shown in D).

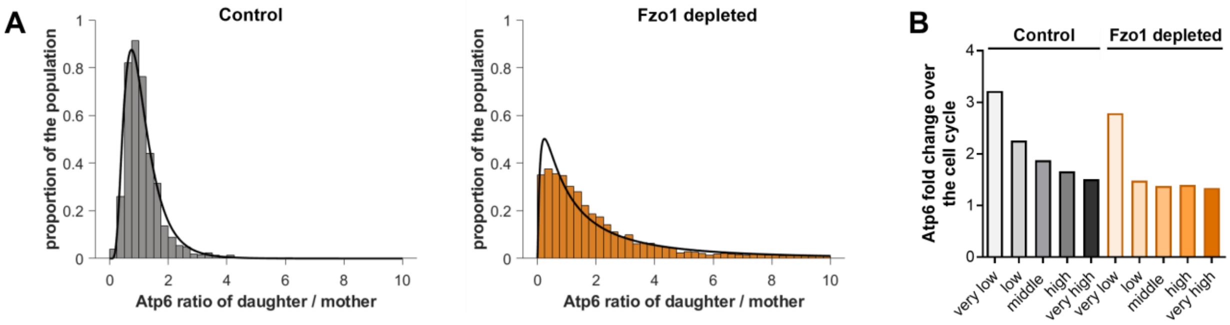


**Supplementary Figure 7:** A) Histogram and lognormal fit (lines) of the ratio of Atp6 between daughters and their mothers at the first time cells are detected in G1 after division is finished. B) average fold-change of Atp6 over the cell cycle of Control and Fzo1 depleted cells with different Atp6 content of the mother at the beginning of the cell cycle. mtDNA synthesis was determined based on the Atp6 content at the beginning of G1. See methods for further details on the categorization.

**Table S1. Strains used in this study.** All strains are W303 derivatives. All strains were constructed in this study.

| Name | Genotype | Description | Figures |
|---|---|---|---|
| LD137-1 | *Mat a, leu2-3,112:: TetR-LEU, can1-100 ura3-1::URA3-Tet-pr-OsTIR1F74G, his3-11,15::His3* | Tetracycline inducible TIR, based on (Azizoğlu et al., 2023) | Fig S1D |
| LD138-1 | *Mat a, LEU, can1-100 ura3-1::URA, his3-11,15::HIS3* | Wild-type | Fig S1D, 2H, S2J |
| LD162-1 | *Mat a, leu2-3,112:: TetR-LEU, can1-100 ura3-1::URA3-Tet-pr-OsTIR1F74G, his3-11,15::HIS3, FLAG-AID-Fzo1* | Tetracycline inducible TIR, FLAG-AID degron (Yesbolatova et al., 2020) | Fig 1A, 1D, S1A, S1B,C, S1D, 2A,B,D,G,H, S2A,B,C,D,F,G, H,I,K |
| LD146-1 | *Mat a, LEU, can1-100 ura3-1::URA, his3-11,15::HIS3, Fzo1::HygMX* | Fzo1 deletion | Fig 1D, 2H, S2D,H,I,J |
| LD181-1 | *Mat a, LEU, can1-100 ura3-1::URA, his3-11,15::His3, 3xFLAG-AID-Fzo1* | FLAG-AID degron | Fig 1A, S1D |
| LD219-1 | *Mat a, leu2-3,112:: TetR-LEU, can1-100 ura3-1::URA3-Tet-pr-OsTIR1F74G, his3-11,15::HIS3, FLAG-AID-Fzo1, Pdr5::HphMX* | Tetracycline inducible TIR, FLAG-AID, Pdr5 deletion | Fig 2C, S2E, |
| LD188-1 | *Mat a, leu2-3,112:: TetR-LEU, can1-100 ura3-1::URA3-Tet-pr-OsTIR1F74G, his3-11,15::TEF-pr-preSu9-mCardinal-His5* | Tetracycline inducible TIR | Fig S1B,C, |
| LD190-1 | *Mat a, leu2-3,112:: TetR-LEU, can1-100 ura3-1::URA3-Tet-pr-OsTIR1F74G, his3-11,15::TEF-pr-preSu9-mCardinal-His5, FLAG-AID-Fzo1* | Tetracycline inducible TIR, FLAG-AID degron, preSu9-mCardinal | Fig S1A, S3G, 4A,B,C,D,E,F,H,I S4A,B,E |
| LD195-1 | *Mat a, leu2-3,112:: LEU2, can1-100 ura3-1::URA3, his3-11,15::TEF-pr-preSu9-mCardinal-His5, Fzo1::HygMX* | Fzo1 deletion, preSu9-mCardinal | Fig S1B,C, 4A, S4A,B |
| LD206-1 | *Mat a, leu2-3,112:: LEU2, can1-100 ura3-1::LexA-ER-AD-LexA-Pr-OsTIR1F74G-Ura3, his3-11,15::TEF-* | Estradiol inducible TIR, FLAG-AID, MitoLoc reporter to | Fig 3A,C,D, S3C,D,E,F. 4G, S4C,D |

| | | | |
|---|---|---|---|
| | *pr-preSu9-mCardinal-His5, 3x-FLAG-AID-Fzo1, TEFpr-preCox4-NG-HO-homology-KanMX* | estimate protein import changes, modified from (Vowinckel et al., 2015) | |
| KK124-1 | *Mat a, leu2-3,112:: LEU2, can1-100 ura3-1::URA3, his3-11,15::TEF-pr-preSu9-mCardinal-His5, TEFpr-preCox4-NG-HO-homology-KanMX, Fzo1::HygMX* | Fzo1 deletion, MitoLoc reporter to estimate protein import changes | Fig S3E |
| LD208-1 | *Mat alpha, ade2-1 ::Ade2, his3-11,15::TEF-pr-preSu9-mCardinal-His5, trp1-1::Trp1, leu2-3,112::Leu2, ura3-1::LexA-ER-AD-LexA-Pr-OsTIR1F74G-Ura3, can1-100, ATP6-NG, 3xFLAG-AID-Fzo1* | Atp6-NG (Jakubke et al., 2021), Su9-mCardinal, Estradiol inducible TIR, FLAG-AID degron | Fig 1B,C, S3A,B S4F, 5A,B,C, S5A,B, 6A-D, S6A-F. S7A,B |
| LD209-1 | *Mat alpha, ade2-1 ::Ade2, his3-11,15::TEF-pr-preSu9-mCardinal-His5, trp1-1::Trp1, leu2-3,112::Leu2, ura3-1::LexA-ER-AD-LexA-Pr-OsTIR1F74G-Ura3, can1-100, ATP6-NG, 3xFLAG-AID-Fzo1, Atg32::HphMX* | Atp6-NG, Su9-mCardinal, Estradiol inducible TIR, FLAG-AID degron, Atg32 deletion | Fig S5C,D |
| LD222-1 | *Mat a, leu2-3,112:: TetR-LEU, can1-100 ura3-1::URA3-Tet-pr-OsTIR1F74G, his3-11,15::HIS3, FLAG-AID-Fzo1, Mrps5-3xFLAG-HphMX, Mrps5-Pr-Neongreen-FLAG-KanMX* | Tetracycline inducible TIR, FLAG-AID, Mrps5-3xFLAG, Mrps5-Promoter-reporter | Fig 2E,F |
| LD224-1 | *Mat a, leu2-3,112:: TetR-LEU, can1-100 ura3-1::URA3-Tet-pr-OsTIR1F74G, his3-11,15::HIS3, FLAG-AID-Fzo1, Qcr7-3xFLAG-HphMX, Qcr7-Pr-Neongreen-FLAG-KanMX* | Tetracycline inducible TIR, FLAG-AID, Qcr7-3xFLAG, Qcr7-Promoter-reporter | Fig 2E,F |

**Table S2. Components of 1x Synthetic complete (SC).**

| component | g/L | [mM] final |
|---|---|---|
| Adenine | 0.031 | 0.228 |
| L-Arg (HCl) | 0.021 | 0.098 |
| L-Aspartic acid | 0.103 | 0.773 |
| L-Glutamic acid | 0.081 | 0.548 |
| L-His | 0.021 | 0.133 |
| L-Leu | 0.123 | 0.941 |
| L-Lys (HCl) | 0.031 | 0.169 |
| L-Met | 0.021 | 0.138 |
| L-Phe | 0.051 | 0.311 |
| L-Ser | 0.386 | 3.670 |
| L-Thr | 0.206 | 1.727 |
| L-Tyr | 0.031 | 0.170 |
| L-Trp | 0.041 | 0.201 |
| L-Val | 0.154 | 1.317 |
| Uracil | 0.021 | 0.184 |

**Table S3. Plasmids used in this study.**

| Name | Description | Origin |
|---|---|---|
| pLHG003 | Tet-Promoter-OsTIRF74G-URA3 | This study |
| FRP2061 | Estradiol inducible Cas9-KanMX | (Azizoğlu et al., 2023) |
| FRP2100 | Nat helper plasmid | (Azizoğlu et al., 2023) |
| FRP2370 | Tet-repressor-LEU2 | (Azizoğlu et al., 2023) |
| pLD036 | TEFpr-preSu9-mCardinal-His5 | This study |
| pLD038 | TEFpr-preCox4-Neongreen-HO homology | This study |
| pLD039 | LexA transcriotion factor with Estradiol-Promoter-OsTIRF74G-URA3 | This study |
| pLD043 | Atp17-Promoter-Neongreen-FLAG-HO-homology-KanMX | This study |
| pLD044 | Mrps5-Promoter-Neongreen-FLAG-HO-homology-KanMX | This study |
| pLD046 | Qcr7-Promoter-Neongreen-FLAG-HO-homology-KanMX | This study |

**TableS4. Optical filters for epifluorescence microscopy.**

Filters used for imaging on a Nikon Ti2-E epifluorescence microscope. All described filters are manufactured by Chroma and purchased from AHF.

| Fluorophore | LED Wave-length | Filter Set | Excitation Filter | Dichroic | Emission Filter |
|---|---|---|---|---|---|
| Neongreen | 513 nm | YFP ET Filter Set | ET500/20x | T515lp, Di 25 mm x 36 mm | ET535/30m |
| mCardinal | 575 nm | 585/29 BrightLine HC<br>650/60 BrightLine HC<br>Beamsplitter T610 LPXR | ET585/29x | T610 LPXR, Di 25 mm x 36 mm | ET650/60 |

**TableS5. Exposure times and intensities for epifluorescence microscopy.**

| Fluorophore | Imaged protein | Intensity | Exposure time |
|---|---|---|---|
| mCardinal | preSu9 | 5 % | 100 ms |
| mNeongreen | preCox4 | 10 % | 200 ms |
| mNeongreen | Atp6 | 10 % | 300 ms |

**Table S6. Primers used for DNA-qPCR.** All Primers were purchased from Integrated DNA Technologies (IDT).

| Name | Sequence |
|---|---|
| COX1_fw | CAACGGGGACAATAGCATGC |
| COX1_rev | CGGACAGTTCTTACCTTGCG |
| COB1_fw | AAATTGGAGCATGCCATGTA |
| COB1_rev | AGCGATTTGTCCCATTAAGA |
| COX2_fw | GTTGATGCTACTCCTGGTAGATT |
| COX2_rev | TTGCATGACCTGTCCCACAC |
| ACT1_fw | CACCCTGTTCTTTTGACTGA |
| ACT1_rev | CGTAGAAGGCTGGAACGTTG |

**Supplementary References**


Azizoğlu, A., Loureiro, C., Venetz, J., & Brent, R. (2023). Autorepression-Based Conditional Gene Expression System in Yeast for Variation-Suppressed Control of Protein Dosage. *Curr Protoc*, *3*(1), e647. https://doi.org/10.1002/cpz1.647

Jakubke, C., Roussou, R., Maiser, A., Schug, C., Thoma, F., Bunk, D., Hörl, D., Leonhardt, H., Walter, P., Klecker, T., & Osman, C. (2021). Cristae-dependent quality control of the mitochondrial genome. *Science Advances*, *7*(36), eabi8886. https://doi.org/doi:10.1126/sciadv.abi8886

Vowinckel, J., Hartl, J., Butler, R., & Ralser, M. (2015). MitoLoc: A method for the simultaneous quantification of mitochondrial network morphology and membrane potential in single cells. *Mitochondrion*, *24*, 77-86. https://doi.org/10.1016/j.mito.2015.07.001

Yesbolatova, A., Saito, Y., Kitamoto, N., Makino-Itou, H., Ajima, R., Nakano, R., Nakaoka, H., Fukui, K., Gamo, K., Tominari, Y., Takeuchi, H., Saga, Y., Hayashi, K. I., & Kanemaki, M. T. (2020). The auxin-inducible degron 2 technology provides sharp degradation control in yeast, mammalian cells, and mice. *Nat Commun*, *11*(1), 5701. https://doi.org/10.1038/s41467-020-19532-z
